# Massively parallel genomic perturbations with multi-target CRISPR reveal new insights on Cas9 activity and DNA damage responses at endogenous sites

**DOI:** 10.1101/2022.01.18.476836

**Authors:** Roger S. Zou, Alberto Marin-Gonzalez, Yang Liu, Hans B. Liu, Leo Shen, Rachel Dveirin, Jay X. J. Luo, Reza Kalhor, Taekjip Ha

**Author notes:** Equal contribution (Roger S. Zou, Alberto Marin-Gonzalez).

## Abstract

We present an approach that combines a Cas9 that simultaneously targets hundreds of epigenetically diverse endogenous genomic sites with high-throughput sequencing technologies to measure Cas9 dynamics and cellular responses at scale. This massive multiplexing of CRISPR is enabled by means of novel multi-target gRNAs (mgRNAs), degenerate gRNAs that direct Cas9 to a pre-determined number of well-mapped sites. mgRNAs uncovered generalizable insights into Cas9 binding and cleavage, discovering rapid post-cleavage Cas9 departure and repair factor loading at PAM-proximal genomic DNA. Moreover, by bypassing confounding effects from gRNA sequence, mgRNAs unveiled that Cas9 binding is enhanced at chromatin-accessible regions, and Cas9 cleavage is more efficient near transcribed regions. Combined with light-mediated activation and deactivation of Cas9 activity, mgRNAs further enabled high-throughput study of the cellular response to double strand breaks with high temporal resolution, discovering the presence, extent (under 2 kb), and kinetics (~ 0.5 hr) of reversible DNA damage-induced chromatin decompaction. Altogether, this work establishes mgRNAs as a generalizable platform for multiplexing CRISPR and advances our understanding of intracellular Cas9 activity and the DNA damage response at endogenous loci.

## MAIN

CRISPR-Cas nucleases, and *Streptococcus pyogenes* Cas9 in particular, have revolutionized biomedicine through effective genome manipulation inside living cells^1^. For genome editing, Cas9 first binds to DNA whose sequence is complementary to the guide RNA (gRNA), then induces a double strand break (DSB), initiating a series of DNA damage responses (DDR) that repair and, ideally, modify the DNA sequence in that region^2^. Although several works have shed light on different stages of this process — e.g. identifying Cas9 binding and cleavage sites along the genome^3–5^, unraveling the pathways that repair Cas9 cleavage^6, 7^, or assessing and predicting mutation outcomes of the genome editing process^8, 9^ — many aspects of the intracellular Cas9 behavior and the ensuing DDR remain incompletely characterized. For instance, how Cas9 departs from genomic DNA after cleavage is unclear^10–12^.

Mismatches impair Cas9 binding and cleavage^3^ and some genomic regions are associated with higher editing efficiencies^7, 13, 14^, yet, how genomic context combines with mismatch levels to dictate Cas9 binding and cleavage rates remains unclear. The cellular response to Cas9-induced DNA damage is also complex and warrants further study^6, 15, 16^. In particular, how damage response factors interact with genomic DNA after Cas9 cleavage, and how chromatin affects and is affected by such interactions is still poorly understood^6, 15, 17^.

Better understanding of these CRISPR-associated intracellular processes would mature existing CRISPR technologies, and potentially inspire new tools and applications^16, 18–20^. However, current methods to study these processes have been limited to a few target sites, *in vitro* measurements, reporter systems, or expressed libraries of gRNAs. Studying a few target positions preclude extracting generalizable conclusions or exploring heterogeneity at different genomic locations^21, 22^ while *in vitro* measurements fail to capture the complex chromatin context in which genome editing occurs and do not always generalize inside cells^11–13^. Reporter systems may not reflect endogenous phenotypes^8, 9^; and expressed gRNA libraries introduce variability arising from differences in binding and transcription efficiency between individual gRNAs and poor temporal control, thus obscuring readouts on relative Cas9 activity at different on-target sites and hindering measurements on the kinetics of genome editing events^4, 8^.

Here, we present an approach whereby a single gRNA, that we denote multi-target gRNA (mgRNA), directs Cas9 to simultaneously target over a hundred endogenous positions genome-wide that are well-mapped by high-throughput short-read sequencing. This multi-target CRISPR system enabled us to interrogate the rapid kinetics of Cas9 activity at endogenous sites and the ensuing DNA damage responses with unparalleled throughput. In particular, mgRNAs showed that, after cleavage, Cas9 has stronger binding to the PAM-distal region, while initial recruitment of DSB repair factors happens more efficiently on the PAM-proximal side. In addition, multi-target CRISPR revealed that the same protospacer sequence, when located in different genomic locations, is bound and cleaved by Cas9 with very different efficiencies; and that these differences can be largely accounted for by differences in the chromatin context of the target site. Finally, by combining mgRNAs with light-mediated CRISPR activation and deactivation, we showed that Cas9 cleavage rapidly induces (~30 min) decompaction of chromatin around the cut site, and that nucleosomes are restored to their original pre-cleavage locations shortly (within 1h) after Cas9-induced damage ends. Together, these results establish multi-target CRISPR as a generalizable platform for massively parallel targeted genomic perturbations and advance our understanding of CRISPR-based genome manipulation and cellular DNA damage and repair.

## RESULTS

### Design, discovery, and validation of multi-target gRNAs

To discover Cas9 gRNA sequences with multiple target positions in the genome, we searched for 20 base pair (bp) sequences adjacent to a Cas9 protospacer adjacent motif (PAM) in the human genome with up to three mismatches from a 280 bp short interspersed nuclear element (SINE)^23^. Over 40,000 20 bp sequences were found, each targeting between 2 to over 1000 putative on-target sites **(Fig. 1b)**. The target sites are located throughout the genome, exhibit balanced representation between gene bodies and intergenic regions, and represent multiple epigenetic states^24^ **(Fig. 1c-d, S1a)**.

**Fig. 1:**
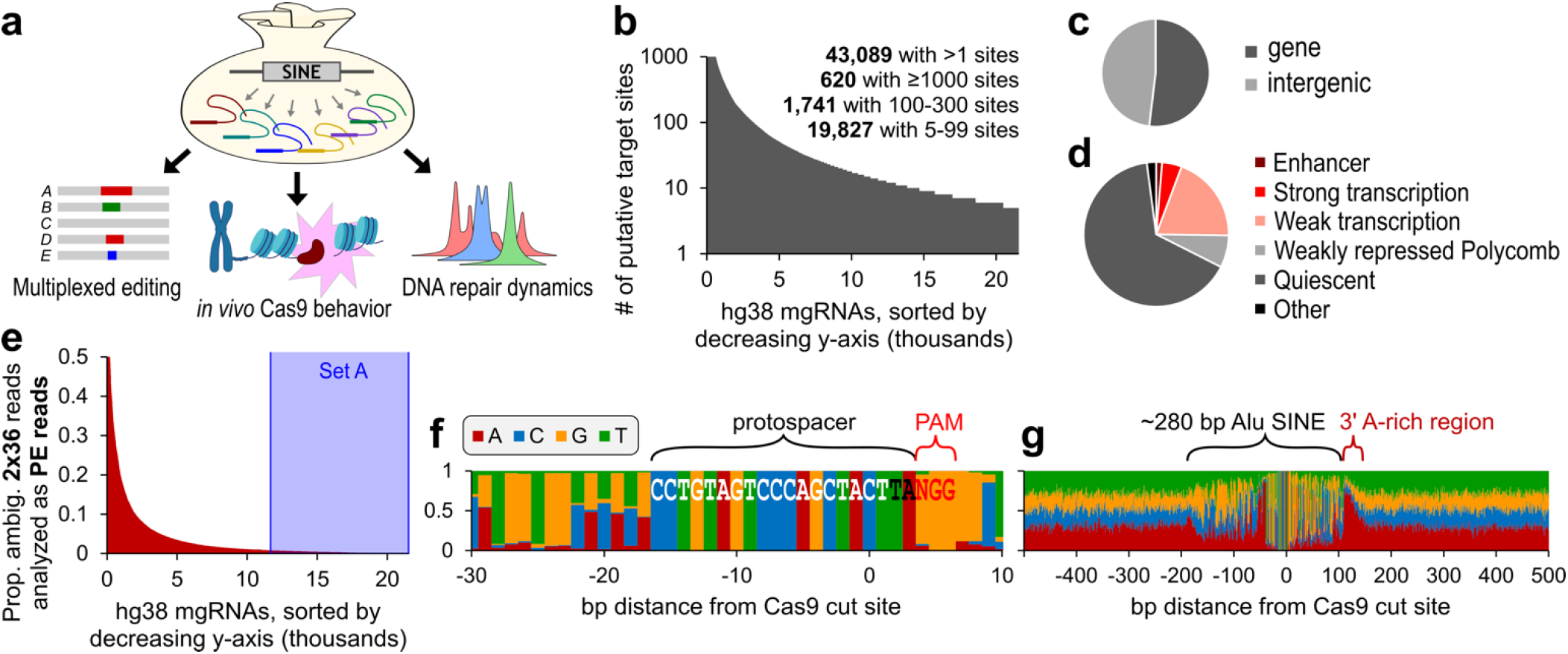
*in silico* characterization of multi-target gRNAs. **a,** Schematic of multi-target gRNA discovery and applications. **b,** Plot of the thousands of unique target sequences identified in silico, sorted along the x-axis by the # of putative on-target sites in the hg38 human genome (y-axis). A maximum of 1000 on-target sites were evaluated for each gRNA (even if the total was higher) to maintain efficient computation. **c,** Approximately half of the target sequences resided in regions annotated as a RefSeq gene. **d,** Distribution of ChromHMM epigenetic labeling for target sequences. Only labels with ≥ 1% representation are directly displayed. Those with < 1% representation are grouped under ‘Other’. **e,** Using bowtie2, we determined the proportion of ambiguous reads from simulated PE 2×36 bp ChIP-seq reads at on-target sites for each target sequence. Target sequences (x-axis) are sorted by decreasing proportion of ambiguous reads (y-axis). Sequences highlighted in blue correspond to Set A, i.e., the set of gRNAs that each have under 1% of ambiguous read alignments from 2×36 bp sequencing. **f-g,** The nucleotide composition at each position around all Cas9 cut sites (A – red, C – blue, G – yellow, T – green) for a select mgRNA. The x-axis labels the base pair distance from the Cas9 cut site at x = 0 (The fourth nucleotide from PAM). (f) Plot displaying a 40 bp window, allowing visualization of the protospacer + PAM sequences that are labeled directly on the graph. The gRNA sequence is written in white and black font. The PAM is written in red font. (g) Plot displaying a 1 kb window. The approximately 280 bp SINE and its 3’ A-rich region are annotated.

Because mgRNAs are derived from short repetitive sequences, we evaluated whether the targeted regions can be uniquely distinguished with high-throughput short-read sequencing. We generated simulated Illumina-style paired-end (PE) 2×36 bp reads at all target sites for each gRNA that have between 5 and 300 target sites. These simulated DNA fragments had a length of between 200 and 600 bp (uniformly distributed); and were randomly chosen to either span the cut site, reside PAM-distal to the cut, or PAM proximal to the cut (see Methods section). We then determined the number of genome-wide alignments for each read using bowtie2^25^. For the majority of gRNAs, only a small minority of reads had ambiguous alignments, i.e., more than one alignment with the same “best” bowtie2 alignment score **(Fig. 1e)**. As expected, treating the PE 2×36 bp reads as single-end (SE) reads increased the percentage of reads with ambiguous alignments **(Fig. S1b)**, whereas increasing the number of sequenced base pairs to 75 at each end (i.e., PE 2×75 bp) reduced this percentage **(Fig. S1c-d)**.

To understand why the surrounding sequences can be unambiguously mapped for most target sites, we aligned the sequence around each expected on-target site for the subset of gRNAs with under 1% ambiguous alignments. The nucleotide composition at each position in a 40 bp window confirmed the expected Cas9 protospacer and showed some sequence diversity immediately outside the protospacer but within the repetitive element **(Fig. 1f)**. Expanding to a 1 kb window confirmed features of the Alu SINE, such as its 280 bp approximate length and A-rich 3’ end^23^ **(Fig. 1g)**. The sequences beyond 150-200 bp from the cut sites were evenly distributed between the four nucleotides and likely correspond to regions that can be uniquely mapped by sequencing. Paired-end sequencing reads can therefore be uniquely mapped given the sequence diversity even within the short repetitive element and the high probability of at least one DNA end being positioned outside the element.

We replicated the same analysis using different genomes (mouse mm10; zebrafish danRer11) and different SINES (rodent B4; zebrafish DR-1 and DR-2)^23^ **(Fig. S1e-o)**. Together, our computational pipeline robustly identified candidate mgRNAs across different genomes, with high degrees of flexibility in both the number of expected target sites and the uniqueness of their flanking sequences.

### Experimental validation of mgRNAs with diverse readouts and cell types

We first experimentally validated the activity of mgRNAs by testing the ability to read genome editing outcomes (mostly insertions and deletions, or indels) at mgRNA targeted sites. HeLa cells were genetically encoded with doxycycline inducible Cas9 and a mgRNA with 10 predicted genome-wide on-target positions. Exposure to doxycycline over the course of ten days followed by nested, multiplexed PCR and amplicon Illumina sequencing at all target positions revealed robust indel generation at 8 out of 10 expected target sites **(Fig. 2a)**. Without doxycycline, we did not observe appreciable levels of mutations **(Fig. S1p)**. Characterization of mutation identity at all target positions revealed diverse mutations but predominantly one nucleotide insertions, consistent with the repair profiles of Cas9-generated DSBs^8, 9, 26, 27^ **(Fig. 2b)**. Similar results were obtained when using a different mgRNA with 20 predicted on-target sites **(Fig. S1q-r)**. Mutation outcome distributions showed high reproducibility between biological replicates **(Fig. 2c, S1s)**. Together, these results demonstrate efficient intracellular mgRNA activity and highly multiplexed analysis at expected target sites using sequencing.

**Fig. 2:**
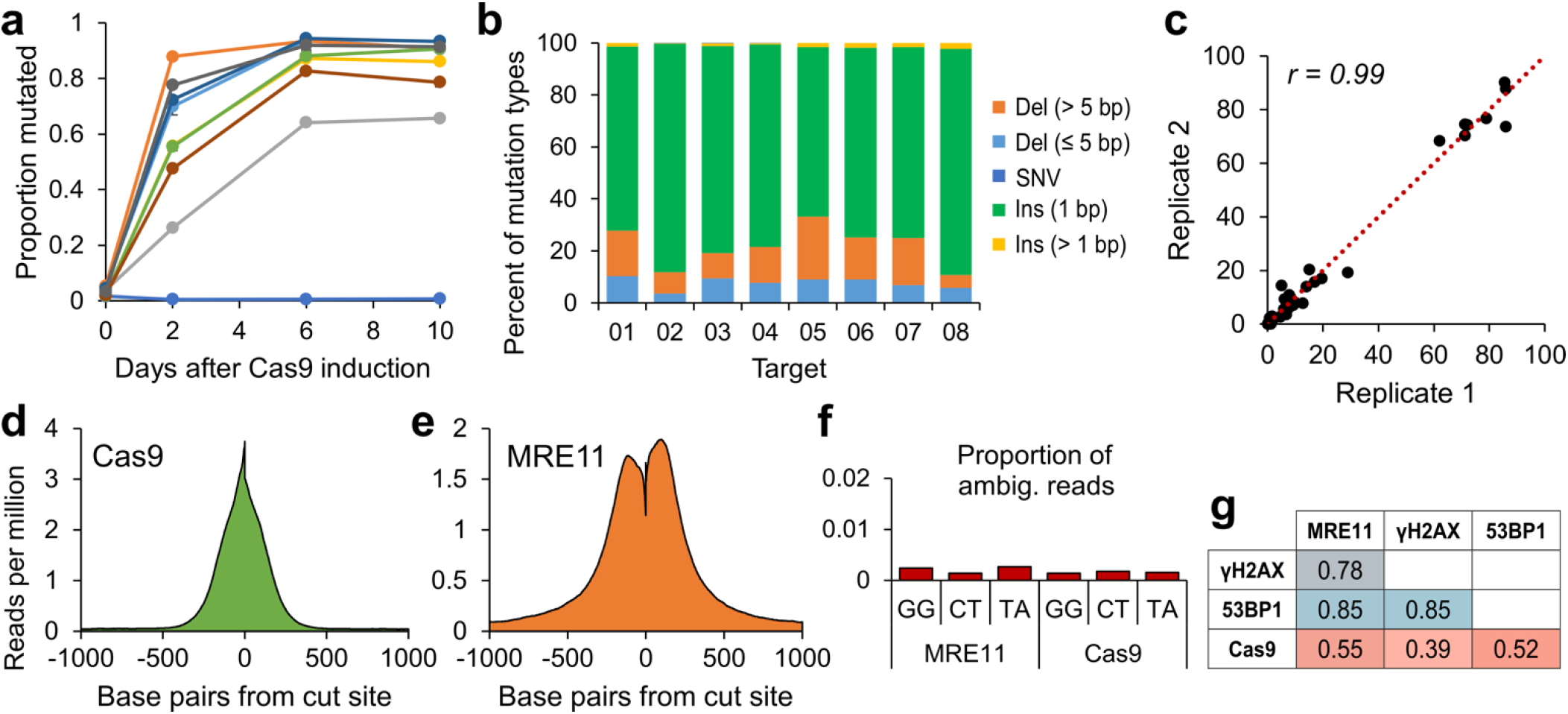
Initial experimental characterization of multi-target gRNAs. **a,** Mutation rate of 10-target mgRNA. HeLa cells with dox-inducible Cas9 were transduced with a 10-target mgRNA and grown in presence of doxycycline. Cells were harvested at different time points (0, 2, 6 and 10 days). The genomic DNA was extracted, PCR amplified and sequenced at the Cas9 target sites (see Methods for details). Data points are the average of 2 biological replicates. **b,** Day-10 samples from Fig. 2a were analyzed to determine the mutational signatures resulting from Cas9 cleavage at each target (8/10) that showed detectable indels. Mutations were classified into five categories: long deletions (longer than 5 bp), short deletions (5 bp or less), single-nucleotide variants (i.e., base change), 1 bp insertions, and insertions longer than 1 bp. Only the mutated protospacers were considered for the analysis. Data points are the average of 2 biological replicates. **c,** Reproducibility analysis of Fig. 2b. Each point represents the percentage of a particular mutation type for a given target (akin to the bars in Fig. 2b, but with replicates analyzed separately). Data points were fitted to a linear function (dotted line) and their Pearson correlation coefficient was computed, yielding r = 0.99. **d-e,** Average profiles of (d) Cas9 and (e) MRE11 enrichment in a 2000 bp window centered at the cut site, across all on-target sites. **f,** Using bowtie2, we determined the proportion of ambiguous reads from measured MRE11 and Cas9 ChIP-seq reads with the 3 target sequences ‘GG’, ‘CT’, and ‘TA’. Reads at all discovered macs2 peaks were used for analysis, which include both on-target and potential off-target sites. **g,** All possible correlations between MRE11, Cas9, γH2AX, and 53BP1 ChIP-seq enrichment measured in a specific window centered at all on-target sites. A 2500 bp window was used for MRE11, a 1500 bp window was used for Cas9, and 20 kb windows were used for γH2AX and 53BP1.

To interrogate Cas9 binding and recruitment of DNA repair factors in a high throughput manner, we tested three mgRNAs (‘CT’, ‘GG’, ‘TA’) with a larger number of on-target sites: 145, 126, and 117, respectively. We delivered a preassembled Cas9-mgRNA RNP into HEK293T cells via electroporation; and 3 hours later we determined the locations of Cas9 binding and cleavage by performing, respectively, chromatin immunoprecipitation with sequencing (ChIP-seq) for Cas9 and for an early DNA damage response protein, MRE11 ^4, 5 15, 28^. ChIP-seq profiles centered and averaged across all on-target sites revealed high enrichment with shapes consistent with previous literature^4, 5, 15, 28^ **(Fig. 2d-e, S2a)**. Cas9 on- and off-target sites were called using Model-based Analysis of ChIP-Seq (MACS2) software^29^, and showed a very small (less than 0.3%) percentage of sequencing reads with ambiguous alignments **(Fig. 2f)**, verifying that ChIP-seq accurately quantified enrichment at sites targeted by these mgRNAs. MRE11 enrichment was highly correlated between biological replicates **(Fig. S2b)** and with other DNA repair markers such as 53BP1 and phosphorylated H2AX (γH2AX)^30^ **(Fig. 2g, S2c-f)**. In contrast, correlations between Cas9 and DNA repair factors were weaker and dependent on gRNA sequence, pointing towards additional factors that affect Cas9 cleavage besides Cas9 binding efficiency (see below) **(Fig. 2g, S2g)**. MRE11 and Cas9 ChIP-seq after multi-target Cas9 delivery could also be performed in induced pluripotent stem cells (iPSCs) **(Fig. S2h-i)**. ChIP-seq enrichment at cut sites was only moderately correlated between iPSCs and HEK293T cells despite high correlation between biological replicates **(Fig. S2j-m)**. This deviation in Cas9 and MRE11 ChIP-Seq enrichment in iPSCs as compared to HEK293T indicates that cell type specific features play a key role in Cas9 activity (see below). Altogether, these results demonstrate multiplexed Cas9 activity and robust ChIP-Seq readout for Cas9 and DNA repair factors at endogenous sites targeted by mgRNAs.

### Cas9 binding and cleavage mechanics at endogenous loci

How Cas9 interacts with genomic DNA is an understudied but important topic essential for better understanding of Cas9 genome editing^5, 10, 28^. For example, whether and how Cas9 departs from genomic DNA after cleavage has been unclear; RNA polymerase^31^ and the histone chaperone FACT^12^ have both been proposed to evict Cas9 but direct evidence inside cells is lacking. mgRNAs present a unique opportunity to dissect these dynamics in a highly multiplexed fashion while controlling for the target sequence. From HEK293T cells exposed to ‘GG’, ‘CT’, or ‘TA’ mgRNAs for 3 hours, Cas9 and MRE11 ChIP-seq reads were categorized as either spanning or abutting the cut site, corresponding to protein-associated DNA fragments that are either intact or cleaved by Cas9, respectively^15, 28^ **(Fig. S3a)**. MRE11 ChIP-seq reads predominantly abutted the cut sites **(Fig. 3a)**, consistent with MRE11 loading on cleaved DNA^15, 28^, whereas Cas9 ChIP-seq reads predominantly spanned the cut sites **(Fig. 3b)**, consistent with Cas9 residing on the target before the cut and departing quickly thereafter. Of the reads that abut each cut site, MRE11 exhibited enrichment bias on the PAM-proximal while Cas9 showed enrichment on the PAM-distal sides of the cut, with the extent of PAM-proximal/PAM-distal bias inversely correlated between MRE11 and Cas9 **(Fig. 3c-f, S3b-f)**. These results suggest stable Cas9 binding before cleavage, ostensibly to check for sequence complementarity, followed by cleavage and rapid release of DNA preferentially from the PAM-proximal side, facilitating MRE11 loading. Therefore, pre-cleavage Cas9 is the dominant DNA bound population, suggesting that cleavage, and not Cas9 dissociation, is hindered or rate-limiting inside cells. Preferential Cas9 dissociation at the PAM-proximal side was observed previously, but only for a single target sequence and only in vitro^32^. Our results validate this observation *in vivo* and further suggest that Cas9 binding to a cleaved DNA terminus can obfuscate it from MRE11 and the cellular DNA damage response.

**Fig. 3:**
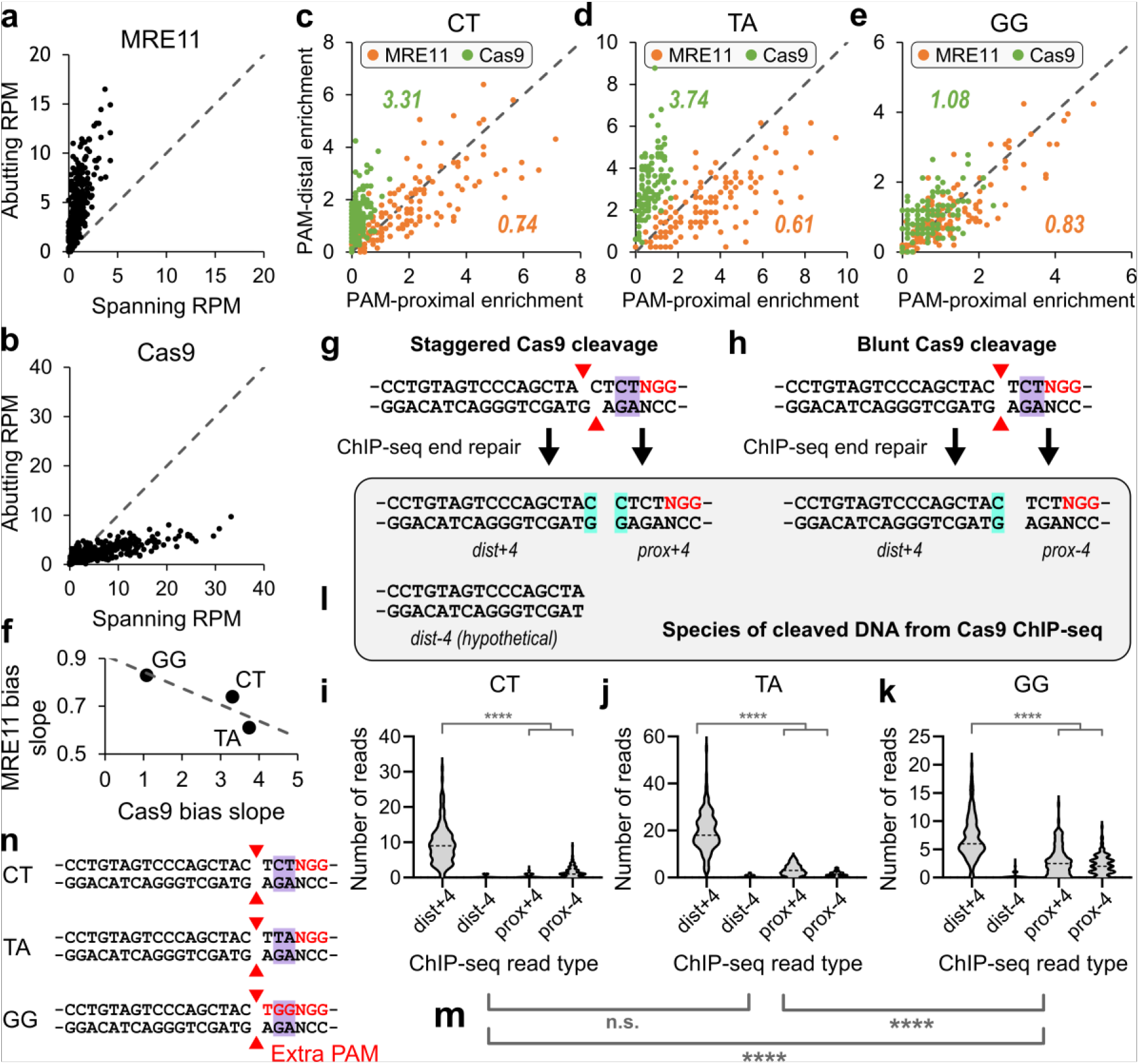
Multiplexed analysis of Cas9 binding and cleavage mechanics revealed by mgRNAs. **a-b,** For each target site, we plot the number of PE reads (per million total sequencing reads) that either span or abut the cut site for (a) MRE11 and (b) Cas9 ChIP-seq. **c-e,** For each on-target site of the (c) ‘CT’, (d) ‘TA’, and (e) ‘GG’ multi-target gRNA sequences, we plot the number of reads (in reads per million) on the PAM-proximal versus PAM-distal side for MRE11 (orange) versus Cas9 (green) ChIP-seq at 3 hours after Cas9 delivery. The orange and green numbers in the graph indicate linear regression slopes of MRE11 and Cas9 enrichment, respectively. **f,** MRE11 and Cas9 slopes from Fig. 3c-e are inversely related as a function of target sequence. **g-h,** Schematic of Cas9 cleavage possibilities and the subsequent ChIP-seq read species. The ‘CT’ target sequence is represented (‘CT’ as the last two nucleotides before PAM, highlighted in purple). Two possibilities for Cas9 cleavage (staggered versus blunt) are displayed, with red triangles annotating the cleavage position at each DNA strand. ChIP-seq end-repair fills in nucleotides at the 3’ end. Only 3 read species are expected based on this model (*dist+4, prox+4*, and *prox-4*). The fourth nucleotide from PAM in all read species is highlighted in cyan. **i-k,** Violin plot of the number of reads, for each Cas9 target site, categorized by four read types (*dist+4, dist-4, prox+4, prox-4*) predicted from models of Cas9 cleavage and ChIP-seq library preparation in Fig. 3g-h. Read species: [*dist+4*: immediately PAM-distal, containing fourth nucleotide from PAM (+4 nucleotide)], [*prox+4*: immediately PAM-proximal containing +4 nucleotide], [*prox-4*: immediately PAM-proximal lacking +4 nucleotide], [*dist-4*: immediately PAM-distal lacking +4 nucleotide – hypothetical]. **l,** *dist-4* is not expected to exist based on our model for Cas9 cleavage and ChIP-seq end repair in Fig. 3g-h. **m,** Student’s t-test of significance for the number of PAM-proximal reads (*prox+4* + *prox-4*) between different gRNA sequences. **n,** Schematic of the ‘CT’, ‘TA’, and ‘GG’ sequences, which only differ in the most PAM-proximal two nucleotides (highlighted in purple). The ‘GG’ target sequence is notable for at least one extra PAM. The ‘NGG’ PAM(s) are labeled in red. The blunt-end cleavage possibility is displayed with red triangles annotating the cleavage position at each DNA strand. **i-k,m,** n.s. indicates not significant, **** indicates p<0.0001.

To further characterize post-cleavage Cas9 mechanics, we modeled Cas9 ChIP-seq read species derived from DNA fragments bound to Cas9 after either staggered or blunt cleavage^33^. From staggered cleavage, DNA end repair during ChIP-seq library preparation fills in the 3’ end, resulting in presence of the fourth nucleotide (from PAM) at both sides of the cut **(Fig. 3g)**. We refer to these ChIP-seq reads on the PAM-proximal and PAM-distal sides as *prox+4’* and *‘dist+4*, respectively. In contrast, from blunt-end cleavage, only the PAM-distal read contains the fourth nucleotide (from PAM), i.e., *‘dist+4’*, whereas the PAM-proximal read does not, resulting in the new *prox-4* read species **(Fig. 3h)**. *‘dist+4’* was significantly more enriched than the sum of *‘prox+4’* and *‘prox-4’* (p < 1E-15, Student’s t-test), recapitulating clear PAM-distal binding bias **(Fig. 3i-l)**. These results suggest that the 16-17 bp of gRNA to genomic DNA base pairing interactions at the PAM-distal side of the cut are stronger than the 3-4 bp of base pairing and PAM-Cas9 interactions at the PAM-proximal side. More importantly, they indicate that the primary mechanism of Cas9 dissociation from DNA is not an inverse of its binding mechanism; rather, an entirely independent pathway which likely involves a distinct set of conformational changes. Interestingly, Cas9 with the ‘GG’ gRNA exhibited significantly stronger association to the PAM-proximal side compared to the other two gRNAs (p < 1E-18) **(Fig. 3m)**, potentially due to the additional ‘NGG’ PAM sequence in the first three nucleotides of the protospacer **(Fig. 3n)**.

### Linking Cas9 binding and DNA damage responses to local epigenetic states

Genome editing efficiencies are difficult to predict but are likely influenced by both sequence and epigenetic factors^3, 7, 13, 14^. Epigenetic influences have been challenging to decipher due to confounding effects of gRNA sequence^21^; mgRNAs are uniquely suited for this task through multiplexed analysis of Cas9 activity with a common gRNA sequence in different epigenetic contexts. To characterize Cas9 binding alone, we measured time-resolved occupancy of (cleavage-deficient) dCas9 using ChIP-seq after multi-target dCas9 delivery. Previous dCas9 ChIP-seq studies were limited by lack of kinetic information and use of standard gRNAs with few target sites^4, 5^. For the multi-target ‘GG’ gRNA, we detected over 4000 dCas9 binding sites with up to 12 mismatches **(Fig. 4a)**. To evaluate Cas9-mediated DNA damage, we measured occupancy of MRE11 after delivery of (cleavage-competent) Cas9. MRE11 was only enriched at sites with 2 or fewer mismatches whereas some sites exhibited clear dCas9 binding for up to over 8 mismatches, and both enrichments were higher if the mismatch solely resided in the PAM-distal region (≥12^th^ position, counting from PAM) **(Fig. 4b-c, S4a)**, consistent with known properties of Cas9 binding and cleavage^5, 11, 34^. Interestingly, there was large heterogeneity in both dCas9 and MRE11 enrichment even between identical on-target sequences **(Fig. 4b-c)**, likely stemming from additional factors such as epigenetics.

**Fig. 4:**
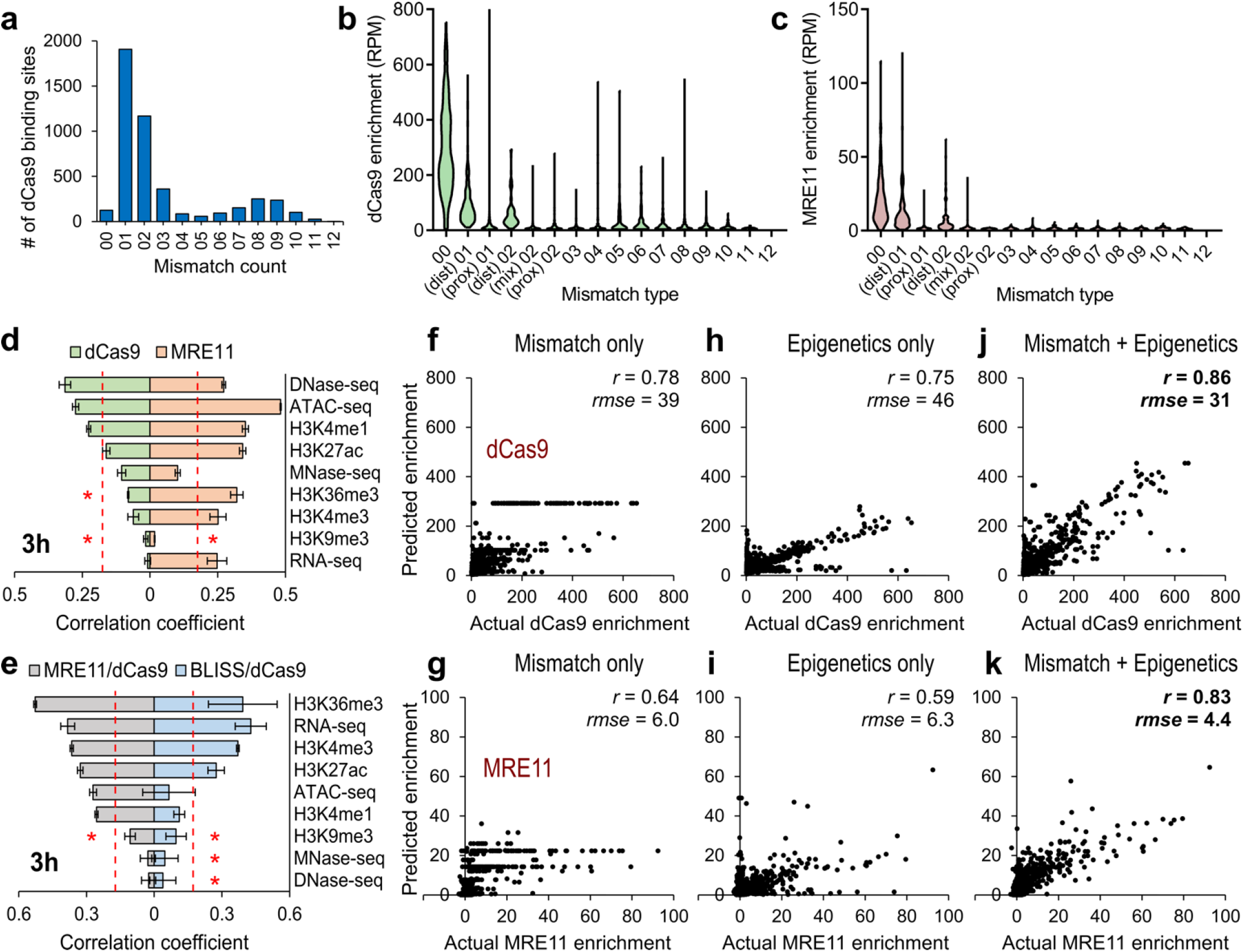
Epigenetic determinants of Cas9 binding and the DNA damage response revealed by mgRNAs. **a,** Identification of all dCas9/’GG’ gRNA binding sites at 3 hours after dCas9 delivery, sorted by mismatch count. ‘00’ corresponds to no mismatches, ‘03’ corresponds to 3 mismatches, and so on. **b,** Histogram of dCas9 enrichment by mismatch type at 3h. ‘(dist) 01’ corresponds to one PAM-distal (≥12^th^ nucleotide, counting from PAM) mismatch, ‘(prox) 01’ corresponds to one PAM-proximal (<12^th^ nucleotide) mismatch, ‘(mix) 02’ corresponds to two total mismatches – one PAM-proximal and the other PAM-distal. **c,** Histogram of MRE11 enrichment at 3 hours after Cas9 delivery, at the same binding positions as evaluated in Fig. 4b. **d,** Absolute value of correlation coefficients between dCas9 (light green) or MRE11 (light orange) enrichment and epigenetic markers at all on-target sites, evaluated at 3 hours after dCas9 or Cas9 delivery, respectively. The order of epigenetic markers from top to bottom is sorted by the degree of its absolute correlation with dCas9 enrichment. The originally negative correlations are marked by red asterisks. **e,** Absolute value of correlation coefficients between dCas9-normalized MRE11 enrichment (MRE11/dCas9; grey) or dCas9-normalized BLISS enrichment (BLISS/dCas9; light blue) with epigenetic markers at all on-target sites, evaluated at 3 hours after (d)Cas9 delivery. The order of epigenetic markers from top to bottom is sorted by the degree of its absolute correlation with MRE11/dCas9 enrichment. The originally negative correlations are marked by red asterisks. **d-e,** Error bars correspond to ±SD from 2 biological replicates. The dotted red lines at 0.173 indicate a significance level cutoff of 0.05. **f-g,** Actual (f) dCas9 and (g) MRE11 enrichment (x-axis) versus predicted enrichment (y-axis) by a trained Random Forest Regressor on an independent test set. Samples evaluated at 3 hours after (d)Cas9 delivery. **h-I,** Same conditions as Fig. 4f-g, except only epigenetic features (RNA-seq, MNase-seq, ATAC-seq, DNase-seq, and H3K4me1, H3K4me3, H3K9me3, H3K27ac, H3K36me3 ChIP-seq enrichment surrounding each cut site) were used as features. **j-k,** Same conditions as Fig. 4f-g, except both mismatch information (Fig. 4f-g) and epigenetic features (Fig. 4h-i) were used as features.

To infer the epigenetic state, we obtained 9 publicly available genome-wide epigenetic maps from the same cell line^35^ and determined their enrichments in specified windows centered around each Cas9 target site **(Fig. S4b)**. dCas9 enrichment was most strongly correlated with markers of DNA accessibility (ATAC-seq and DNase-seq), consistent with prior reports^4, 5^, whereas MRE11 recruitment was correlated with many other epigenetic features **(Fig. 4d)**, suggesting additional epigenetic factors are at play beyond Cas9 binding. To characterize the MRE11 damage response independent of Cas9 binding, we normalized MRE11 signal by dCas9 signal at all cut sites, which yielded the strongest correlation with gene bodies (H3K36me3 and RNA-seq), promoters (H3K4me3), and enhancers (H3K27ac) **(Fig. 4e)**. This suggests either higher Cas9 cleavage efficiencies or more efficient MRE11 recruitment at these genomic regions, which we can distinguish by directly measuring DSB levels genome-wide using Breaks Labeling *In Situ* and Sequencing (BLISS)^36^. BLISS enrichment was highly correlated with MRE11 (*r* = 0.7) **(Fig. S4c-d)**, and the pattern of epigenetic correlation for dCas9-normalized BLISS enrichment (unrepaired DSBs given the same amount of Cas9 binding) mirrored dCas9-normalized MRE11 enrichment **(Fig. 4e)**. These results suggest that identical Cas9 on-target sites bound by the Cas9:gRNA complex are cleaved at different rates. While the reason(s) for this behavior is unclear at this time, regions near gene bodies, promoters, and enhancers exhibit intrinsically higher cleavage activity by a bound Cas9, which, along with better Cas9 binding near accessible chromatin, likely contribute to the previously observed higher editing efficiencies in these regions^3, 5, 13, 14^.

### Prediction of genome editing processes using machine learning

To further explore the determinants of Cas9 binding and DNA damage induction, we trained Random Forest machine learning models to predict both dCas9 and MRE11 enrichment at all binding locations. From solely mismatch information, dCas9 and MRE11 enrichment at 3 hours could be adequately predicted for an independent test dataset with r = 0.78 and 0.64, respectively **(Fig. 4f-g)**. Using solely epigenetic information led to comparable levels of performance with r = 0.75 for dCas9 and 0.59 for MRE11 **(Fig. 4h-i)**. However, using both mismatch and epigenetic information greatly improved prediction, resulting in r = 0.86 for dCas9 and 0.83 for MRE11 **(Fig. 4j-k)**. Comparable levels of predictive power were also achieved for the early 30 min time point **(Fig. S4e-j)**. These results highlight the importance of local epigenetic state in modulating Cas9 activity and provides further evidence that combining epigenetic with mismatch information improves the prediction of genome editing activity^13, 37^.

### Kilobase-scale increase in chromatin accessibility at Cas9-induced DNA damage sites

It has been proposed that local chromatin decompaction occurs after DNA damage to facilitate repair, but direct evidence has not been observed at single Cas9 DSBs^38, 39^. To measure chromatin accessibility changes after DNA damage, we performed Assay for Transposase-Accessible Chromatin with sequencing (ATAC-seq)^40^ with and without exposure to multi-target Cas9. Averaged background-subtracted ATAC-seq enrichment centered at Cas9 target sites exhibited locally increased accessibility after 3 hours of Cas9 exposure **(Fig. 5a-b)**. Excess chromatin accessibility was only detected within 1-2 kb from the cut site (p < 9E-5) **(Fig. 5c)**. The average full width at half maximum (FWHM) of ATAC-seq chromatin accessibility increase was similar to but greater than that of MRE11 (722 bp versus 523 bp, respectively). **(Fig. S5a-c)**. Furthermore, there was no clear correlation between MRE11-normalized ATAC-seq enrichment and any epigenetic marker **(Fig. S5d)**, suggesting that chromatin opening after Cas9 cleavage occurs independent of chromatin context.

**Fig. 5:**
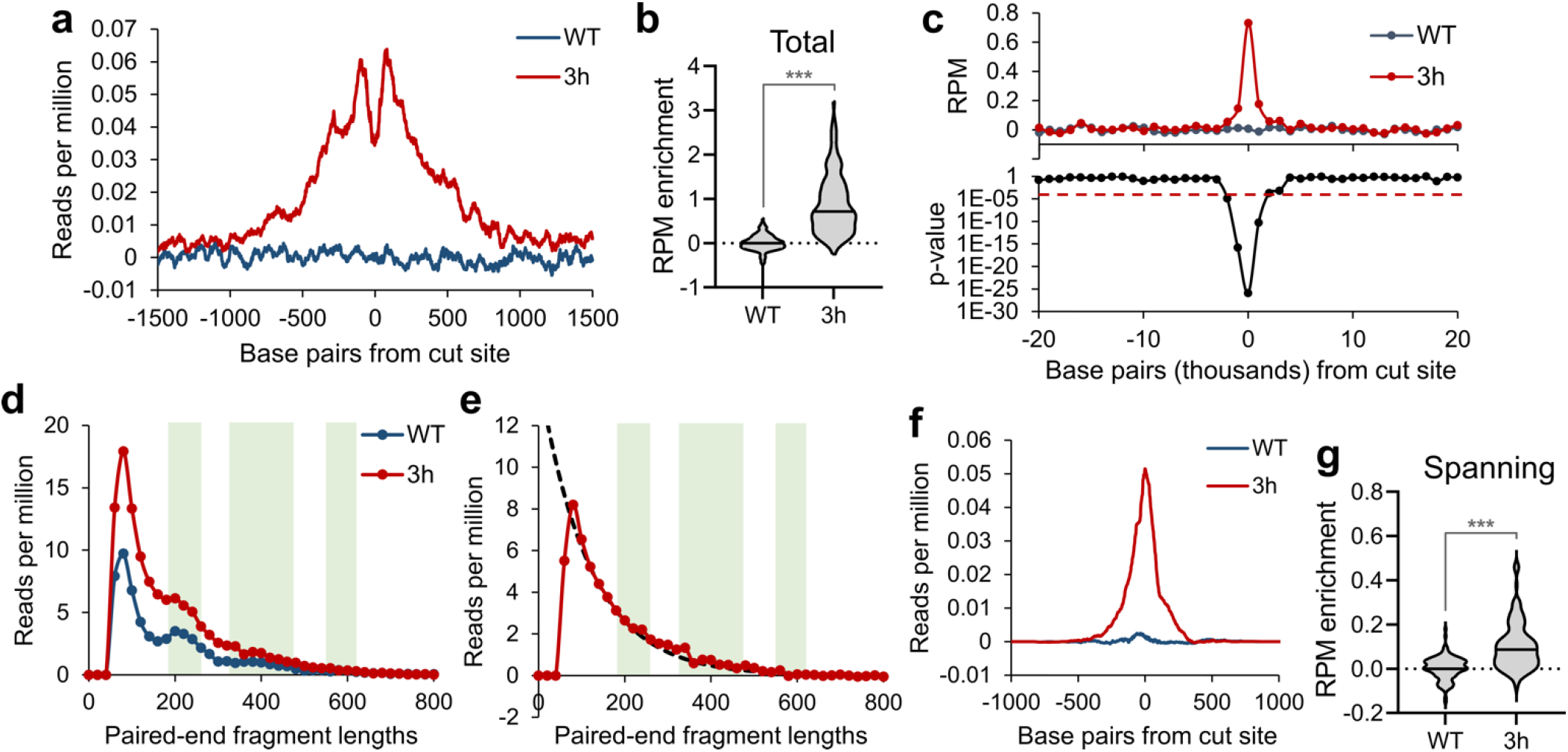
Study of chromatin accessibility change after DNA damage using mgRNAs. **a,** Averaged background-subtracted ATAC-seq profiles across all on-target sites for cells without Cas9 (WT, blue), or 3 hours after Cas9/‘GG’-gRNA delivery (3h, red). ‘Background-subtracted’ enrichment was obtained by subtracting the number of reads at each position for Cas9-negative cells from Cas9-exposed cells, yielding enrichment values that quantify ‘excess’ accessibility due to Cas9 exposure. **b,** Violin plots of background-subtracted ATAC-seq enrichment (in reads per million, RPM) at each target site, for cells without Cas9 (WT), or 3 hours after Cas9/GG’-gRNA delivery (3h). *** indicates p<0.001. **c,** (top) Average background-subtracted ATAC-seq enrichment (RPM) in 1 kb windows moving upstream and downstream of all cut sites, for cells without Cas9 (WT), or 3 hours after Cas9/GG’-gRNA delivery (3h). (bottom) Student’s t-test p-values of difference in enrichment between ‘WT’ and ‘3h’ samples in each 1 kb window. **d,** Histogram of ATAC-seq sequenced DNA lengths for cells without Cas9 (WT, blue), or 3 hours after Cas9/GG’-gRNA delivery (3h, red). **e,** Subtraction of the ‘3h’ sample by ‘WT’ sample from Fig. 5g. Fitting (black dotted curve) was performed using an exponential decay model. **f-g,** Same as Fig. 5a-b, but only for the subset of ATAC-seq reads that span the cut site.

Next, we inferred the lengths of all paired-end ATAC-seq reads around expected target sites. For cells without Cas9, the distribution of sequencing read lengths within a window of 1.5 kb from the target site showed a local maximum that corresponded to nucleosome occupancy footprinting^40^ **(Fig. 5d, S5e)**. Cells exposed to Cas9 had excess ATAC-seq reads, and we determined the length distribution of the excess reads **(Fig. 5e, S5f)**. The resulting distribution lacked the nucleosomal footprinting signature and was well-fit by an exponential decay, consistent with distances between adjacent Tn5 transposition events that are assumed to be a Poisson point process **(Fig. 5e, S5f)**. Assuming nucleosome spacing length of around 200 bp, this implies that the up to 2 kb accessible region **(Fig. 5c)** lost up to 10 nucleosomes^41, 42^. We further uncovered a sub-population of ATAC-seq reads spanning the target sites that significantly increased after Cas9 delivery **(Fig. 5f-g, S5g-h)**, which must correspond to post-cleavage DNA that has undergone ligation and suggests that chromatin recompaction does not occur immediately after ligation. In conclusion, Cas9 cleavage induces a localized, nucleosome-depleted, kilobase-scale region of increased accessibility that can persist after DNA ligation, which potentially facilitates the binding of DNA damage-associated proteins such as repair factors, cohesin, and transcription factors to promote successful repair^43 44 45^.

### Light-responsive mgRNAs reveal the timescales of DSB repair and chromatin accessibility changes

The temporal sequence of events after Cas9 cleavage has not been well-characterized but can be explored using the very fast light-activatable CRISPR system (vfCRISPR) based on a photocaged gRNA (cgRNA)^15^. We delivered Cas9 with the multi-target ‘GG’ cgRNA to HEK293T cells, waited 12 hours for stable Cas9 binding, then light-activated Cas9 and performed time-resolved BLISS, MRE11 ChIP-seq, and ATAC-seq. DSBs and MRE11 damage responses were undetectable before light activation, confirming that Cas9 is inactive without light exposure **(Fig. 6a-b)**. As early as 10 min after activation, BLISS exhibited the strongest relative enrichment increase followed by MRE11 ChIP-seq signal **(Fig. 6a-b)**, consistent with initial DSB induction followed by repair protein recruitment. ATAC-seq enrichment increased by 30 min after Cas9 activation but not 10 min **(Fig. 6c, S5i)**, suggesting that DSB-induced increase in accessibility occurs downstream of initial repair protein recruitment **(Fig. 6d)**.

**Fig. 6:**
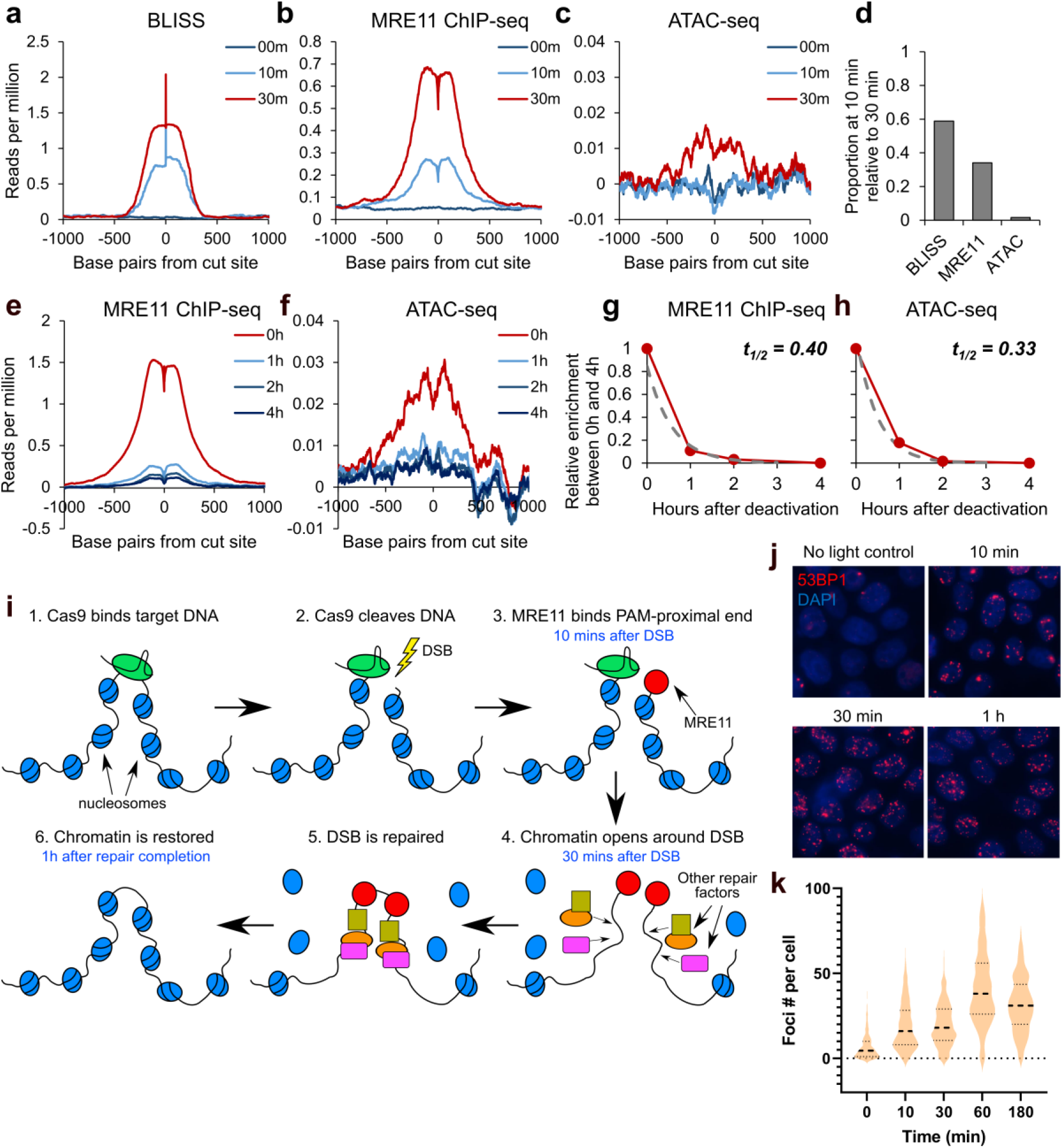
Timescales of DNA damage response recruitment and dissolution revealed by very fast CRISPR activation and deactivation. **a-c,** Average (a) BLISS, (b) MRE11 ChIP-seq, and (c) ATAC-seq enrichment across all on-target sites using Cas9 in complex with very fast CRISPR (vfCRISPR) ‘GG’ gRNA. Samples were evaluated at 00 min (no Cas9 activation), 10 min, and 30 min after Cas9 activation using light. **d,** Quantification of Fig. 6a-c. Proportion, at 10 min, of the maximal enrichment at 30 min after Cas9 activation. **e-f,** Average (e) MRE11 ChIP-seq and (f) ATAC-seq enrichment across all on-target sites using Cas9 in complex with ‘GG’ photocleavable gRNA (pcRNA). Samples were evaluated at 0h (no deactivation), 1h, 2h, and 4h after Cas9 deactivation using light. **g-h,** Quantification of Fig. 6e-f. We plotted the relative change in (g) MRE11 ChIP-seq and (h) ATAC-seq enrichment between 0h and 4h after Cas9 deactivation. **i,** Cartoon summarizing our findings on DNA damage response in the context of Cas9 cleavage. Within ~10 min after Cas9 cleavage, MRE11 is recruited preferentially to the PAM-proximal side. ~30 min after Cas9 cleavage, chromatin undergoes decompaction around the cut site, potentially to facilitate the recruitment of additional DNA repair factors. Once the DSB is repaired, chromatin accessibility returns to the original pre-DSB state. **j,** Representative images of 53BP1 immunofluorescence microscopy as a function of time after ‘GG’ multi-target Cas9 activation in HEK293T cells using the light-inducible vfCRISPR system. Cas9/gRNA was electroporated into cells 12h before light-based Cas9 activation. **k,** Quantification of number of 53BP1 foci in each cell from Fig. 6j at all evaluated time points.

After repair of DNA damage, the duration of accessibility increase remains unknown. However, without an effective method for CRISPR deactivation, intracellular Cas9 will repeatedly cleave repaired loci and preclude measurements of chromatin restoration^46^. We therefore employed a light-deactivatable Cas9 based on a photocleavable gRNA (pcRNA) to synchronize Cas9 deactivation, facilitating chromatin profiling through repair completion^46^. We delivered Cas9 with multi-target ‘GG’ pcRNA to HEK293T cells, deactivated Cas9 after 2 hours, then performed time-resolved MRE11 ChIP-seq and ATAC-seq. After Cas9 deactivation, MRE11 enrichment rapidly declined across all target sites with a mean half-life of 0.4 hours (**Fig. 6e)**, which likely corresponds to completion of DNA repair^15^. There was no detectable correlation between MRE11 departure and the tested epigenetic markers **(Fig. S5j)**. Chromatin accessibility also declined with a comparable mean half-life of 0.33 hours **(Fig. 6f-h)**, demonstrating that damage-induced accessibility reverses at similar timescales as MRE11 departure.

Altogether, our findings on Cas9 activity and DNA damage response are summarized in **Fig. 6i**. After binding and cleavage of target DNA, Cas9 quickly releases the DNA preferentially from the PAM-proximal side, enabling binding of MRE11 to this DNA end. This recruitment of MRE11 occurs within 10 min of DSB generation. Afterwards, within 30 min of DSB, chromatin undergoes decompaction, whereby nucleosomes get evicted in a window of ~1 kb centered at the cut site. This removal of nucleosomes potentially facilitates recruitment of additional DNA repair factors. Once the lesion has been repaired, the nucleosomes are repositioned around the cut site, restoring the chromatin accessibility landscape to pre-cleavage levels.

## DISCUSSION

Here, we reported the discovery and diverse applications of multiplexed CRISPR using multi-target gRNAs. We first identified and characterized tens of thousands mgRNAs that each target 2 to over 1000 positions in the human, mouse, and zebrafish genomes, providing an extensive and immediately available resource for rapid adoption of mgRNAs for diverse applications. We then combined mgRNAs with high-throughput sequencing readouts to provide the most comprehensive study to date of genome editing processes at endogenous loci **(Table S1)**. These findings advance our understanding of CRISPR genome editing and DNA damage responses in numerous ways. First, the large number and diversity of target sites enable generalizable observations, from discovering the destabilizing impact of even one PAM-distal mismatch by quantification of Cas9 binding at thousands of sites, to characterizing the extent of local chromatin decompaction after DNA damage via ATAC-seq at hundreds of cut sites. Second, comparison between different target sites reveals site-specific heterogeneity, such as better Cas9 binding at chromatin accessible regions and better cleavage by bound Cas9 near transcribed regions. Third, aggregating data across multiple target sites boosts readout signal and enables new observations, such as aggregating ATAC-seq reads across all target sites to measure local nucleosome depletion after Cas9 DNA damage. Fourth, the compatibility of mgRNAs with very fast CRISPR activation and deactivation^15, 46^, allowed us to quantify with high temporal resolution the dynamics of chromatin accessibility during the repair of Cas9-mediated DNA damage. Cas9 with mgRNAs also exhibits several advantages over “multi-target” meganucleases^38^, including programmable target positions, precise time control using CRISPR activation and deactivation, facile delivery without the need to generate a stable cell line, and direct relevance to CRISPR genome editing. Finally, the ability to read mutational outcomes of multi-target CRISPR via next-generation sequencing paves the way towards its use as a new system for endogenous genetic barcoding, by leveraging the indels introduced by Cas9 at endogenous sites as cellular barcodes. Supporting this claim, the indels at 8 target sites generated by the 10 target mgRNA in HeLa cells with doxycycline inducible Cas9 demonstrated high barcoding diversity, as measured by Shannon entropy^47^ **(Fig. S6a)**.

One potential concern of mgRNA is the relatively high number of simultaneous DSBs in each cell. These DSBs can average under 50 per cell for a 126-targeting mgRNA based on the number of 53BP1 foci in immunofluorescence microscopy **(Fig. 6j-k)** and have variable effects on cell division. However, it is important to note that this effect can be largely controlled by using mgRNAs with a proper number of targets **(Fig. S6b)**. In our case, we primarily used mgRNAs with over 100 target sites to increase multiplexing and because all experiments were conducted within a short period of time (3 hours or less within Cas9 delivery) during which no altered cellular phenotypes were observed **(Fig. S6c-d)**. In addition, at such timescales, the relative kinetics of Cas9 binding, Cas9 cleavage and repair events at different target sites are likely unaffected by the high mutation load. Nevertheless, for other applications of mgRNAs, the appropriate number of on-target sites should be determined.

Multi-target CRISPR using mgRNAs presents a powerful avenue for multiplexed genomic manipulation at endogenous loci. The discoveries enabled by mgRNAs have broad implications for CRISPR-based genome manipulation and DNA repair studies. We expect that mgRNAs can be adapted to study other CRISPR-Cas systems that are being discovered or engineered, such as base editors and prime editors^20, 48, 49^. Multiplexed Cas9 targeting may also be applied to DNA repair studies using live-cell imaging, where the nature of DNA damage is well-defined unlike in studies using radiation or chemical mutagens^44, 50^. mgRNAs may also be useful as a genetic barcoding tool, due to their ability to introduce mutations at endogenous sites, thus eliminating the need for reporter systems^33, 47, 51^. Finally, inducible expression of gRNAs targeting millions of sites may act as a cell death switch to alleviate adverse effects of therapeutic cell therapy^52, 53^. We envision multi-target CRISPR using mgRNAs as a versatile platform that furthers the development and understanding of broadly applicable genome manipulation technologies.

## Acknowledgements

We thank Dr. Sophie Z. Gu for detailed edits of the manuscript. We thank Prof. Geraldine Seydoux for access to the Lonza 4D nucleofector system. We thank the Johns Hopkins Transcriptomics and Deep Sequencing Core for Illumina sequencing.

## Author contributions

R.S.Z. and T.H. conceived the project. R.S.Z., A.M.G designed and performed experiments. R.S.Z analyzed data and wrote the manuscript with contributions from all authors. Y.L., and H.B.L. assisted with ChIP-seq. L.S. and R.D. assisted with cell line generation and cloning. J.X.J.L assisted with data analysis. R.K. assisted with the design of mutation kinetics experiments. T.H. supervised the project.

## Data and materials availability

All data associated with this study are present in the paper or Supplementary Materials. All sequencing data is uploaded to Sequence Read Archive under BioProject accession PRJNA733683 with analysis code available on GitHub (https://github.com/rogerzou/multitargetCRISPR).

## Methods

### SpCas9 purification

BL21-CodonPlus (DE3)-RIL competent cells (Agilent Technologies 230245) were transformed with Cas9 plasmid (Addgene #67881), then inoculated into 5 mL of LB media with Ampicillin at 1 μL/mL concentration. The bacteria culture was grown overnight (37 °C, 220 rpm), then transferred to 1 L of LB-Ampicillin media supplemented with 0.1% glucose until OD_600_ of ~0.5. The cells were then induced with 0.2 mM IPTG and maintained overnight at 18 °C. The bacteria were then pelleted at 4500 × *g*, 4 °C for 15 min and resuspended in 20 mL of lysis buffer containing 20 mM Tris pH 8.0, 250 mM KCl, 20 mM imidazole, 10% glycerol, 1 mM TCEP, 1 mM PMSF, and cOmplete™ EDTA-free protease inhibitor tablet (Sigma-Aldrich 11836170001). This cell suspension was lysed using a microfluidizer and the supernatant containing Cas9 protein was clarified by spinning down cell debris at 16,000 × *g*, 4 °C for 40 min, then filtering with a 0.2 μm syringe filter (Thermo Scientific™ F25006). Ni-NTA agarose bead slurry (Qiagen 30210) was pre-equilibrated with 5 column volumes of lysis buffer. The clarified supernatant was then loaded at 4 °C, and protein-bound Ni-NTA beads were washed with 15 column volumes of wash buffer containing 20 mM Tris pH 8.0, 800 mM KCl, 20 mM imidazole, 10% glycerol, and 1 mM TCEP. Gradient elution was performed with buffer containing 20 mM HEPES pH 8.0, 500 mM KCl, 10% glycerol, and varying concentrations of imidazole (100, 150, 200, and 250 mM) at 7 mL collection volume per fraction. The eluted fractions were evaluated on an SDS-PAGE gel and imaged with Coomassie blue (Bio-Rad 1610400) staining. To remove potential DNA contamination, 1 mL Q Sepharose^®^ column (GE Healthcare 17051005) was charged with 1 M KCl and then equilibrated with elution buffer containing 250 mM imidazole. The purified protein solution was then passed over the Q column at 4 °C. The flow-through was collected and dialyzed in a 10 kDa SnakeSkin™ dialysis tubing (Thermo Fisher Scientific 68100) against 2 L of 20 mM HEPES pH 7.5, and 500 mM KCl, 20% glycerol at 4 °C, overnight. The next day, the protein was dialyzed for an additional 3 hours in fresh dialysis buffer. The final Cas9 protein was concentrated to 10 μg/μL using Amicon^®^ Ultra 10 kDa centrifugal filter unit (Millipore UFC801024), aliquoted, flash-frozen, then stored at −80 °C.

### Cell culture

HEK293T cells (ATCC^®^ CRL-3216™) and HeLa (ATCC^®^ CCL-2™) cells were cultured at 37 °C under 5% CO_2_ in Dulbecco’s Modified Eagle’s Medium (DMEM, Corning) supplemented with 10% FBS (Clontech), 100 units/mL penicillin, and 100 μg/mL streptomycin (DMEM complete). Cells were tested every month for mycoplasma.

A human induced pluripotent stem cell (hiPSC), WTC11 cell line^54^, was used for all iPS cell experiments in this study. We followed the guidelines of Johns Hopkins Medical Institute for the use of this hiPSC line. Briefly, frozen WTC11 cells were first thawed in 37 °C water bath and washed in Essential 8 Medium (E8; Thermo Fisher Scientific, #A1517001) by centrifugation. After resuspension, WTC cells were plated onto a 6 cm cell culture dish pre-coated with human embryonic cell (hES cell)-qualified matrigel (1:100 dilution, Corning #354277). Plate coating should be performed for at least 2 h. Subsequently, 10 μM ROCK inhibitor (Y-27632; STEMCELL, #72308) was supplemented into the E8 medium to promote cell growth and survival. For subculture, WTC11 cells were dissociated from the plate using accutase (Sigma, #A6964) and passaged every 2 days. WTC11 cells were maintained in an incubator at 37 °C with 5% CO_2_.

### Measurements of mutations at mgRNA targets

#### Dox-inducible Cas9 cell line

A PiggyBac system was used to transpose HeLa cells (ATCC^®^ CCL-2™) with a vector carrying Cas9 under the control of a Tet-On inducible promoter and a puromycin resistance gene. Two days after transposition, clonal cell lines were isolated and grown in presence of 2 μg/mL of puromycin.

#### Plasmid and lentivirus production for transduction of mgRNA

mgRNAs were purchased as forward and reverse ssDNA oligos on IDT DNA, containing, respectively, a 5’-CACCG, and a 5’-CAAA and 3’-C overhang (see Supplemental Table 5 for mgRNA oligos sequences). Vectors carrying 10-target or 20-target mgRNAs were made by cloning mgRNAs into the LentiGuide-Hygro plasmid (Addgene #139462). In short, the LentiGuide-Hygro plasmid was digested using BsmBI-v2 enzyme (NEB #R0739), and the larger band (~9 kb) was gel extracted. Forward and reverse oligos were annealed and phosphorylated, and then ligated overnight into the digested vector. E. coli cells (NEB # C2987) were transformed with the ligation product and plated following manufacturer’s instructions. The following day, individual colonies were selected and grown in selection media; and plasmids were purified the next day using QIAprep Spin Miniprep Kit (Qiagen, #27106). Correct insertion of the mgRNA, and other key features of the final plasmid, were verified via Sanger sequencing.

For lentivirus production, Lenti-X™ 293T cells (takarabio #632180) were grown in 10 cm dishes up to ~70% confluency. 5.25 μg of transfer plasmid (LentiGuide-Hygro with 10-target or 20-target mgRNA) was mixed with 0.75 μg of pMD2.G (Addgene #12259) and 1 μg of psPAX2 (Addgene #12260), and with 21 μL of TransIT-Lenti (Mirus #6603). The mixture was incubated for ~15 min and then added dropwise to the 10 cm dish containing Lenti-X cells. The viral supernatant was collected at 36 h, 48 h, and 60 h; and was then filtered and concentrated using Lenti-X™ Concentrator (takarabio #631232), according to the manufacturer’s instructions.

#### Transduction of dox-inducible Cas9 cells with lentivirus carrying mgRNA plasmids

Dox-inducible Cas9 monoclonal cells were grown to ~60 % confluency in 6-well plates. Cells were exposed to virus carrying mgRNA (~0.3-0.5 MOI) and 8 μg /mL polybrene for 24 h. Two days after infection, cells were exposed to 100 μg /mL hygromycin and were kept under such selection conditions for all subsequent experiments. Death of half of the cells confirmed successful integration of the plasmid at the estimated MOI.

#### Induction of Cas9 for mutation of mgRNA targets

For each mgRNA, an initial set of cells genomically integrated with dox-inducible Cas9 and mgRNA were harvested prior to doxycycline addition for the zero-time point. Cells were then grown in 24-well plates in exposure to 2 μg/mL of doxycycline to induce Cas9 expression. At different time points after doxycycline induction, a number of cells were harvested at the time of passaging and were subjected to gDNA extraction. For the 10-target mgRNA, a second experiment was performed in parallel where cells were grown and harvested under the same conditions, but in absence of doxycycline.

#### Sequencing of mgRNA targets

In order to measure mutations of mgRNA targets, gDNA was extracted from Cas9-mgRNA cells using Qiagen DNeasy Blood & Tissue kit (Qiagen #69506). The final product was eluted in 60 uL of elution buffer and quantified using QuBit (Thermo). gDNA was then subjected to a total of three PCRs: two nested PCRs to amplify the target region and a third, indexing PCR to attach the NGS adapters and indices. 1 ng of input DNA was put into the first PCR, or PCR-1, which was run to 20 cycles using the primers shown in Supplemental Table 6. 1 μL of 1:10 dilution of unpurified PCR-1 product was used for the second nested PCR. PCR-2 was run to 20 cycles, using the primers shown in Supplementary Table 7. The product from PCR-2 was purified using 1x volume of AMPure XP beads (Beckman Coulter) and eluted in 15 μL of IDTE buffer (IDT DNA). 1 μL of this purified product was used for PCR-3, which was run to 7 cycles using the primers from Supplementary Table 8. The final product was purified using 0.8x volume of AMPure XP beads, eluted in 15 μL of IDTE, and quantified using QuBit. Products from different samples were pooled and sequenced using a MiSeq (Illumina).

We found conditions for pooling primers from different targets that yielded a balanced representation of all the sequenced targets among the NGS reads. In other words, PCR-1 and PCR-2 were run in a multiplexed fashion. This enabled a considerable saving of time and reagents. For the 10-target mgRNA, we pooled all the PCR-1 primers and all the PCR-2 primers (totaling one set of primers per PCR) in equimolar amounts to a final concentration of 5 μM per oligo. For the 20-target mgRNA, we made three sets of primers per PCR: set-1 with targets 2-6, set-2 with targets 8-11, and set-3 with targets 1, 7 and 12. Targets were then de-multiplexed during the data analysis (see below).

### Determining mutation levels and mutation outcomes of mgRNAs

In order to determine the mutation levels of the different mgRNA targets, we first de-multiplexed these targets (which were amplified in a multiplexed fashion using pooled primer sets). For that, we aligned the first 50 bp of each paired-end read to the human genome. A given read was considered to contain an mgRNA target if the paired-end alignment fell within a window of 1,000 bp from the expected genomic location of the target. A mutation was called if the intact theoretical protospacer sequence was not found in the read.

For classification of the mgRNA target mutations, we first defined, for each target site, two key sequences that were, respectively, 20 bp upstream and downstream of the theoretical genomic location of the Cas9 cut site. For each read that aligning to a target site, these two key sequences were identified and the distance between them was computed. Reads with distances shorter than the expected value were classified as deletions, while reads with distances longer than expected were classified as deletions. Reads with the expected distance between the key sequences, but with mutations in the protospacer were classified as single-nucleotide variants (SNV).

### Electroporation of Cas9 RNP

crRNA and tracrRNA sequences are in Table S2. 2 μL of 100 μM crRNA was mixed with 2 μL of 100 μM tracrRNA (Integrated DNA Technologies) and heated to 95 °C for 5 min in a thermocycler, then allowed to cool on benchtop for 5 min. To form the RNP complex, 3 μL of 10 μg/μL (~66 μM) of purified Cas9 was mixed with the annealed 4 μL 50 μM cr:tracrRNA, then 8 μL of dialysis buffer (20 mM HEPES pH 7.5, and 500 mM KCl, 20% glycerol) was mixed in for a total of 15 μL. This solution was incubated for 20 min at room temperature to allow for RNP formation.

HEK293T cells were maintained to a confluency of ~90% prior to electroporation. 12 million cells were trypsinized with 5 min incubation in the incubator, then 1:1 of DMEM complete was added to inactivate trypsin. This mixture was centrifuged (3 min, 200 × *g*), supernatant removed, followed by resuspension of the cell pellet in 1 mL PBS, centrifugation (3 min, 200 × *g*), and finally complete removal of supernatant. 90 μL of nucleofection solution (16.2 μL of Supplement solution mixed with 73.8 μL of SF solution from SF Cell Line 4D-Nucleofector™ X Kit L) (Lonza) was mixed thoroughly with the cell pellet. The 15 μL RNP solution was mixed in along with 2 μL of Cas9 Electroporation Enhancer (Integrated DNA Technologies). The entirety of the final solution (approximately 125 μL) was transferred to one well of a provided cuvette rated for 100 μL. Electroporation was then performed according to the manufacturer’s instructions on the 4D-Nucleofector™ Core Unit (Lonza) using code CA-189. Some white residue may appear in the cell mixture after electroporation, but that is completely normal. A total of 400 μL of DMEM complete was used to completely transfer the cells out of the cuvette, before plating to culture wells pre-coated with 1:100 collagen. A minimum of 4 million cells are used for each ChIP. For time-resolved experiments, this means one electroporation equates to 3 samples.

For WTC-11 iPSCs, cells were dissociated from the plate using accutase (Sigma, #A6964). Electroporation was performed using the Lonza P3 Primary Cell 4D-Nucleofector™ X Kit L using code CA-137, on 10 million cells, and using 65 μL of the P3 solution mixture with EP enhancer per electroporation cuvette (compared to 90 μL of comparable SF solution mixture for HEK293T cells). After electroporation, cells were resuspended in E8 medium supplemented with 10 μM ROCK inhibitor (Y-27632; STEMCELL, #72308), and plated onto a 10 cm cell culture dish pre-coated with human embryonic cell (hES cell)-qualified matrigel (1:100 dilution, Corning #354277) for at least 2 hours.

### Chromatin immunoprecipitation sequencing

The ChIP protocol was adapted from previous literature^28^. This protocol describes the reagents for one ChIP. When more than one ChIP can be performed, reagent master mixes can be prepared whenever appropriate. Briefly, cells were washed once with room temperature PBS, then washed off the plate with 10 mL DMEM and transferred to 15 mL falcon tubes. 721 μL of 16% formaldehyde (methanol-free) was added for 12 min in room temperature. 750 μL of 2 M glycine was added to quench the formaldehyde. Cells were spun down with 1,200 × *g* at 4 °C for 3 min, then washed with ice-cold PBS twice, spinning down with the same centrifugation conditions. Pellet can be decanted, flash-frozen, then stored in −80 °C for later use. Cells were then resuspended in 4 mL lysis buffer LB1 (50 mM HEPES, 140 mM NaCl, 1 mM EDTA, 10% glycerol, 0.5% Igepal CA-630, 0.25% Triton X-100, pH to 7.5 using KOH, add 1x protease inhibitor right before use) for 10 min at 4 °C, then spun down 2,000 × *g* at 4 °C for 3 min. The supernatant was decanted. Cells were then resuspended in 4 mL LB2 (10 mM Tris-HCl pH 8, 200 mM NaCl, 1 mM EDTA, 0.5 mM EGTA, pH to 8.0 using HCl, add 1x protease inhibitor right before use) for 5 min at 4 °C, spun down with the same protocol, and the supernatant decanted. Cells were then resuspended in 1.5 mL LB3 (10 mM Tris-HCl pH 8, 100 mM NaCl, 1 mM EDTA, 0.5 mM EGTA, 0.1% Na-Deoxycholate, 0.5% N-lauroylsarcosine, pH to 8.0 using HCl, add 1x protease inhibitor right before use) and transferred to 2 mL Eppendorf tubes for sonication with 50% amplitude, 30 s ON, 30 s OFF for 12 min total time (Fisher 150E Sonic Dismembrator). Sample was spun down with 20,000 × *g* at 4 °C for 10 min, and supernatant was transferred to 1.5 mL LB3 in a 15 mL falcon tube. 300 μL of 10% Triton X-100 was added, and the entire solution was well mixed by gentle inversion.

For WTC11 iPSCs, the only difference is a 7 min fixation time versus 12 min for HEK293T cells. Empirically, 7 min reduced overfixing for iPSCs resulting in better MRE11 ChIP signal.

Beads pre-loaded with antibodies were prepared before cell harvesting. 50 μL Protein A beads (Thermo Fisher) were used per IP and transferred to a 2 mL Eppendorf tube on a magnetic stand. Beads were washed twice with blocking buffer BB (0.5% BSA in PBS), then resuspended in 100 μL BB per IP. 3 μL of antibody per IP (Cas9 – Diagenode C15310258; MRE11 – Novus NB100-142; γH2AX – Abcam ab81299; 53BP1 – Novus NB100-305) was added and placed on rotator for 1-2 h. Right before IP, the 2 mL tube was placed on a magnetic rack and washed 3x with BB, before resuspending in 50 μL BB per EP. 50 μL of beads in BB were transferred to each IP and placed in 4 °C rotator for 6+ hours.

Samples were transferred to 2 mL Eppendorf tubes on a magnetic stand, washed 6x with 1 mL RIPA buffer (50 mM HEPES, 500 mM LiCl, 1 mM EDTA, 1% Igepal CA-630, 0.7% Na-Deoxycholate, pH to 7.5 using KOH), then washed 1x with 1 mL TBE buffer (20 mM Tris-HCl pH 7.5, 150 mM NaCl), before decanting. Beads containing ChIP-ed DNA were mixed with 70 μL elution buffer EB (50 mM Tris-HCl pH 8.0, 10 mM EDTA, 1% SDS) and incubated 65 °C for 6+ hours. 40 μL of TE buffer was mixed to dilute the SDS, followed by 2 μL of 20 mg/mL RNaseA (New England BioLabs) for 30 min at 37 °C. 4 μL of 20 mg/mL Proteinase K (New England BioLabs) was added and incubated for 1 h at 55 °C. The genomic DNA was column purified (Qiagen) and eluted in 35 μL nuclease free water.

Oligonucleotide sequences for library preparation are in Table S3. End-repair/A-tailing was performed on 17 μL of ChIPed DNA using NEBNext^®^ Ultra™ II End Repair/dA-Tailing Module (New England BioLabs), followed by ligation (MNase_F/MNase_R) with T4 DNA Ligase (New England BioLabs). 10, 13, 13 and 13 cycles of PCR using PE_i5 and PE_i7XX primer pairs were performed for γH2AX, 53BP1, Cas9, and MRE11 ChIP samples, respectively to amplify libraries. Samples were pooled, quantified with QuBit (Thermo), Bioanalyzer (Agilent) and qPCR (BioRad), then sequenced on a NextSeq 500 (Illumina) using high-output paired 2×36bp reads. Reads were demultiplexed after sequencing using bcl2fastq. Paired-end reads were aligned to hg19 or hg38 using bowtie2. Samtools was used to filter for mapping quality >= 25, remove singleton reads, convert to BAM format, remove potential PCR duplicates, and index reads.

### Genome-wide DSB detection with BLISS

The BLISS protocol was adapted from previous literature^36^. All oligonucleotide sequences are provided in Table S4. BLISS adapters are annealed by mixing 5 μL of Top, 5 μL of Bottom, with 40 μL of nuclease free water (NFW) in a PCR tube, heated to 95 °C for 5 min, then cooled to 4 °C at −0.5 °C per 30 seconds.

#### 5’ Adenylation of RA3

5’ Phosphorylated RA3 oligonucleotides were adenylated using 5’ DNA Adenylation Kit (New England BioLabs E2610S), by mixing 1 μL of 100 μM of 5’ Phosphorylated RA3, 13 μL of NFW, 2 μL of 10x reaction buffer, 2 μL of 1 mM ATP, and 2 μL of Mth RNA ligase to a total 20 μL volume in a PCR tube. Samples were heated in a thermocycler to 65 °C for 1 hour, then 85 °C for 5 minutes, then ethanol precipitated to 10 μL of NFW.

#### Cas9 delivery

Follow protocol from ‘Electroporation of Cas9 RNP’, then plate 400,000 cells each to 24-well.

#### In situ cell lysis

All PBS washes were performed by gently adding 500 μL of 1xPBS to a 24-well at room temperature (RT), incubating at RT, then gently removing all liquid from the well. Cells were washed 1x with PBS, fixed in the 24-well with 4% paraformaldehyde at RT for 10 min, then washed 3x with PBS. Cells were incubated with lysis buffer BLISS-LB1 (10 mM Tris-HCl pH 8, 10 mM NaCl, 1 mM EDTA, 0.2% Triton X-100) for 1 hour at 4 °C, washed with PBS, then incubated with BLISS-LB2 (10 mM Tris-HCl pH 8, 150 mM NaCl, 1 mM EDTA, 0.3% SDS) for 1 hour in a 37 °C incubator. Cells were then washed 2x with PBS.

#### In situ DSB blunting to ligation

Wash 2x with 200 μL of 1x CutSmart Buffer (New England BioLabs) with 2 min incubation RT. Add 150 μL of Quick Blunting Kit (New England BioLabs E1201L) (115.5 μL NFW 15 μL 10x blunting buffer, 1.5 μL 10 mg/mL BSA, 15 μL 1 mM dNTPs, 3 μL Blunt Enzyme mix), incubate 1 hour at RT. Wash 2x with 200 μL of 1x CutSmart Buffer with 2 min incubation at RT. Add 150 μL of dA-Tailing mix (New England BioLabs E6053L) (131 μL NFW, 15 μL of 10x Tailing buffer, 4 μL Klenow Fragment), incubate 30 min at 37°C. Wash 2x with 200 μL of 1x CutSmart Buffer with 2 min incubation RT. Wash with 200 μL 1x T4 Ligase Buffer with 5 min incubation at RT. Add 150 μL of Ligation mixture (New England BioLabs M0202L) (110 μL NFW, 15 μL of 10x T4 Ligase Buffer, 12 μL 10 mM ATP, 7.5 μL 20 mg/mL BSA, 4 μL of 10 μM ligated BLISS adapter, 1.5 μL T4 Ligase). Incubate for 2 hours at RT. To remove unligated adapters, wash 4x with high salt wash (HSW) buffer (10 mM Tris-HCl pH 8, 2 M NaCl, 2 mM EDTA, 0.5% Triton X-100) with 15 min incubation at 37 °C, 1x with PBS at RT (2 min incubation), 1x with NFW at RT (2 min incubation). Then, add 95 μL of DNA extraction buffer (10 mM Tris-HCl pH 8, 100 mM NaCl, 50 mM EDTA, 1% SDS), add 5 μL Proteinase K (New England BioLabs), scrape cells from 24-well, transfer 50 μL each to two PCR tubes, incubate overnight at 55 °C. The next day, purify with spin columns (Qiagen MinElute), elute in 300 μL TE buffer.

#### Sonication

Used sonicator (Qsonica Q125), with 30% amplitude, 10s ON, 10s OFF, 1.5 min total time. Run on 2% gel to verify sonication size to 300-500 bp. Clean up reaction with 0.8x volume AMPure XP beads (Beckman Coulter), elute to 25 μL NFW.

#### In vitro transcription and RA3 adapter ligation

Assemble 20 μL IVT reaction (New England BioLabs – E2050S) (8 μL of template in NFW, 10 μL NTP reaction buffer, 2 μL T7 RNAP mix), incubate at 37 °C for 4 hours in thermocycler. Add 2 μL DNaseI to remove DNA template. Cleanup IVT reaction using Monarch RNA cleanup kit (New England BioLabs – T2040L), eluting in 12 μL NFW. Prepare RA3 adapter ligation with T4 RNA Ligase 2, truncated KQ (New England BioLabs – M0373S) (5 μL IVT RNA product, 1 μL 10x reaction buffer, 2 μL of 50% PEG8000, 1 μL of 5’ adenylated RA3 primer, 0.5 μL of T4 RNA Ligase 2 truncated KQ, 0.5 μL RNaseOUT (Thermo)). Incubate 2 hours in RT, perform RNA cleanup with Monarch RNA cleanup kit, eluting in 12 μL NFW.

#### Reverse transcription and library amplification PCR

Prepare RT reaction with SuperScript III (Thermo Fisher – 18080044). (all 12 μL of RA3 adapter-ligated sample, 1 μL of 10 μM RTP primer, 1 μL of 10mM dNTP mix). Heat to 65 °C for 5 min, then transfer to ice for 1 minute. Add (4 μL 5x First-Strand Buffer, 1 μL 100mM DTT, 1 μL RNaseOUT, 1 μL SuperScript III). Incubate 50 °C for 1 hour, then 70 °C for 15 min, store in −20 °C. For library amplification PCR, add 1 μL of RT sample, 9.5 μL of NFW, 1 μL of 10 mM RP1 primer, 1 μL of 10 mM RP_X primer, 12.5 μL of Q5 2x master mix (New England BioLabs). Cycle on thermocycler: 10s at 98 °C, 15 cycles of 30s at 60 °C and 30s at 72 °C, then 72 °C for 10 min. Perform 0.8x volume AMPure XP cleanup to 20 μL of elution buffer. Run on a 2% agarose gel to remove low molecular weight adapter dimers, isolating for fragments between 300 bp and 1 kb.

#### Sequencing

Samples were pooled, quantified with QuBit (Thermo), Bioanalyzer (Agilent) and qPCR (BioRad), then sequenced on a NextSeq 500 (Illumina) using high-output paired sequencing, with 64 bp for read1 and 36 bp for read2. The reason for this difference is that read1 starts with a 12 bp unique molecular index (UMI), followed by a 13 bp constant adapter region (CGCCATCACGCCT). This leaves 39 bp of genome-specific reads for read1. Read2 still has 36 bp genome-specific reads.

#### Sequencing output processing

Reads were demultiplexed after sequencing using bcl2fastq. Only the subset of reads with the correctly matching 13 bp constant adapter region (CGCCATCACGCCT) in read1 was used for subsequent genome-side alignment. Paired-end reads were aligned to hg19 or hg38 using bowtie2. Samtools was used to filter for mapping quality >= 25, remove singleton reads, convert to BAM format, remove potential PCR duplicates, and index reads.

### ATAC-Seq

In order to measure changes in accessible chromatin associated to DSB repair, 900,000 HEK293T cells were electroporated as described above (Electroporation of Cas9 RNP section), but using reduced reagents’ volumes: 1 μL of crRNA, 1 μL of tracrRNA, 1.5 μL of Cas9, 1.5 μL of dialysis buffer and 20 μL of nucleofection solution, using SF Cell Line 4D-Nucleofector™ X Kit S (32 RCT) (3.6 μL of Supplement solution and 16.4 μL of SF solution). After electroporation, cells were transferred to 3 mL of complete media and plated into two wells of a 12-well plate pre-coated with collagen. In parallel, 400,000 non-electroporated cells were harvested and were plated in another well of the same 12-well plate. Three hours after electroporation, cells were washed with PBS and harvested via scraping. Cells were counted and 50,000 cells were used for ATAC for each condition (electroporation and no-electroporation).

ATAC-Seq was performed following the Omni-ATAC protocol from^55^ with minimal modifications. Harvested cells were centrifuged for 5 min at 500 × *g*, 4 °C in a swinging-bucket rotor. The supernatant was carefully removed so as not to perturb the small, barely visible pellet. Cells were then resuspended in 50 μL of cold lysis buffer (10 mM Tris-HCl pH 7.4, 10 mM NaCl, and 3 mM MgCl_2_ supplemented with 0.1% Igepal-CA630, 0.1% Tween-20, and 0.01% digiton), gently mixed (pipetting up and down three times) and incubated on ice for 3 min. 1 mL of wash buffer (10 mM Tris-HCl pH 7.4, 10 mM NaCl, and 3 mM MgCl_2_ supplemented with 0.1% Tween-20) was then added and was mixed by inverting the tube three times. Nuclei were then centrifuged at 500 × *g* for 10 min, 4 °C in a swing-bucket rotor. The supernatant was carefully discarded and the nuclei were resuspended in 50 μL of transposition reaction (25 μL 2xTD buffer, 16.5 μL PBS, 5 μL NF H_2_O, 1 μL 10% tween-20, 1 μL 1% digitonin, 2.5 μL transposase) and incubated at 37°C for 30 min in a Thermomixer with shaking at 1,000 rpm. After transposition, DNA was purified using a Qiagen MinElute PCR purification kit (elute in 21 μL of elution buffer).

Transposed DNA was amplified using the conditions and primers described in^56^. After the initial pre-amplification step, we found that very few additional cycles are needed (typically two or three) to achieve one third/fourth of the saturation in qPCR. The amplified product was purified using double-sided AMPure bead purification (0.5x and 1.3x) to remove both primers and large products (>1 kb). Final DNA libraries were eluted in 32 μL of IDTE. 5 μL were used in a 2% agarose gel to check for the quality of the library.

For vfCRISPR-ATAC, electroporation was performed as described above (only using caged gRNA). Electroporated cells were split into three wells across several 12-well plates pre-coated with collagen (300,000 cells/well). Cells incubated for 12 h before shining light. UV-light exposure conditions are the same as in (Liu et al., 2020). Time-points (10 min, 30 min) denote time from UV exposure to cell lysis. The rest of the ATAC protocol was performed as described above.

For pcRNA-ATAC, cells were electroporated and split into four wells across four different 12-well plates pre-coated with collagen. Cells incubated for 2 h before shining light.

Samples were pooled, quantified with QuBit (Thermo), Bioanalyzer (Agilent) and qPCR (BioRad), then sequenced on a NovaSeq 500 (Illumina) using paired 2×50 bp reads. Reads were demultiplexed after sequencing using bcl2fastq. Paired-end reads were aligned to hg19 or hg38 using bowtie2. Samtools was used to filter for mapping quality >= 25, remove singleton reads, convert to BAM format, remove potential PCR duplicates, and index reads. Finally, because NovaSeq outputs two lanes for each sample, the two lanes for each sample were merged using samtools.

### CRISPR activation and deactivation

The special cgRNA or pcRNAs were used in the place of normal crRNAs when complexed with tracrRNA. For activation, Cas9/cgRNA was first electroporated into cells, plated onto 12-wells plates, then incubated for 12 hours to allow stable Cas9 binding, but not cleavage. Next, cells were exposed to 1 min of 365 nm light exposure from a handheld blacklight (Amazon https://www.amazon.com/JAXMAN-Ultraviolet-365nm-Detector-Flashlight/dp/B06XW7S1CS/). Either 1, 3, or 6 flashlights were used at once. When multiple flashlights are used, they are conveniently held together using a 3D-printed flashlight holder. (https://github.com/rogerzou/chipseq_pcRNA/blob/master/Jaxman_LED_flashlight_holder_design/files/8zeFEC_PViSo.stl). Samples were harvested without light exposure, or 10m and 30m after light exposure.

For deactivation, Cas9/pcRNA was first electroporated into cells, plated onto 12-well plates, incubated for 2 hours, then exposed to light of the same dose. Samples were harvested during the time of light exposure, or at 1h, 2h, and 4h after light exposure.

### Immunofluorescence microscopy of 53BP1 foci after multi-target Cas9 activation

To check DNA damage upon vfCRISPR induction, we evaluated the number of endogenous 53BP1 foci in cells through immunofluorescence microscopy. Briefly, we electroporated Cas9/Alu RNP into HEK293 cells as previously described. After 1 h incubation, we illuminated the cell samples with 365 nm light for 30 s to trigger Cas9 cleavage activity. The samples were fixed with 4% of paraformaldehyde in 1x PBS for 10 min as a function of time (0, 10 min, 30 min, 1 h and 3 h) and subsequently quenched by 1x PBS supplemented with 0.1 M glycine for 10 min. After thoroughly rinsing with 1x PBS, 0.5% Triton-X was used to permeabilize cell membrane for 10 min. 2% w/v BSA in 1x PBS was used to passivate the sample for 1 h and at room temperature. Without further rinsing, anti-53BP1 antibody (Novus Biological, NB100-304) was diluted in 1:1000 in 1x PBS and directly added into the chamber. After 1 h incubation, primary antibody was removed and the sample was thoroughly washed with 1x PBS three times. Alexa647 conjugated secondary antibody was diluted in 1:1000 and applied to the sample for 1 h. Finally, the sample was rinsed three times and mounted with Prolong Diamond mounting media (Thermo Fisher Scientific) overnight. We imaged all cell samples using Nikon Ti-E fluorescence microscope equipped with Hamamatsu CMOS camera and an objective of 40x magnification. Cell samples were scanned in z-stack with a total depth of 5 micrometers such that all 53BP1 foci (Alexa647) within the cell nuclei (DAPI) were captured. For image analysis, 3D image datasets were first processed into 2D datasets in FIJI using maximum intensity projection. The number of 53BP1 foci per nuclei was automatically analyzed with a custom-built CellProfiler3 pipeline.

### Summary of custom bioinformatics pipeline

We developed companion software available on GitHub (https://github.com/rogerzou/multitargetCRISPR) that is open-source, modular, and well-documented. This software enables in silico discovery and characterization of new multiplexed target sequences along with validation of expected ambiguous sequencing read percentages. After experimental collection of repair factor ChIP-seq, BLISS, and ATAC-seq datasets, an automated pipeline extracts enrichment information and determines relationships between target position, sequencing read enrichment, and local epigenetic state. The software’s modularity enables facile extension to investigate new questions in CRISPR-mediated genome editing.

### Discovery and characterization of multi-target gRNA sequences

Starting from a 280 bp short interspersed nuclear element (SINE) repetitive sequence, for all 20 bp substrings in both the forward the reverse complement direction, obtain all 20 bp sequences with up to 3 mismatches from template restricted to the 9 most PAM-proximal nucleotides. An additional requirement is that these sequences have a GC content of 40-70%. This resulted in 75,626 unique target sequences.

To determine the number of genome-wide alignments for each target sequence, we outputted each gRNA + PAM into a FASTA file, then ran the following command using bowtie2 with ‘-k 1000’ mode, which searches up to 1000 alignments for each line in the FASTA, i.e. each target sequence.

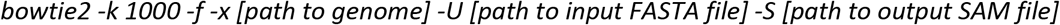

Custom code iterates through all alignments (up to 1000) for each gRNA, then determines whether each alignment is within a RefSeq gene annotation, as well as the ChromHMM epigenetic labeling (Kundaje et al., 2015). Because ChromHMM based on HEK293T cells was not readily available, we curated ChromHMM annotations from A549 (E114), GM12878 (E116), HeLa-S3 (E117), and K562 (E123) cell lines, and the final ChromHMM annotation for each target was the consensus of the four annotations. The annotation data was obtained from https://egg2.wustl.edu/roadmap/web_portal/index.html.

### Ambiguous read proportions from simulated ChIP-seq reads

For gRNA with 100-300 on-target sites in the genome, custom code generated 100 simulated paired-end sequencing reads between 200 to 600 bp in length (taken from uniform distribution). The reads are randomly chosen to either span the cut site, reside PAM-distal to the cut, or PAM-proximal to the cut. For reads PAM-distal or PAM-proximal to the cut, the distance from the edge of the DNA to the cut site samples draw from an exponential distribution. Both 2×36 PE reads and 2×75 PE reads were simulated.

The PE reads were outputted to FASTA files (read1 and read2). bowtie2 was then used to determine up to 10 alignments for each simulated read pair:

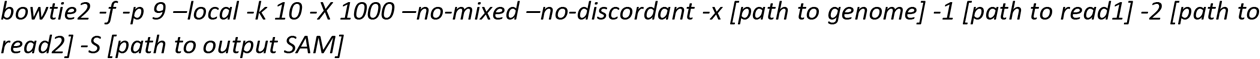

The code subsequently determines whether the original position of the read pairs matches the best alignment based on bowtie2, and whether this best alignment has the uniquely best alignment score (there could be multiple reads with the same best alignment score). The proportion of reads that satisfy these requirements represent the proportion of uniquely best alignments. The proportion of ambiguous alignments is 1 minus this value.

### Ambiguous read proportions from real ChIP-seq reads

Used all dCas9 binding positions for analysis. For each binding position, convert real paired-end ChIP-seq reads found within a specified window width centered at the Cas9 binding site into FASTA read1 and read2 file formats. Then, the protocol follows the previous section, ‘Ambiguous read proportions from simulated ChIP-seq reads’, starting with use of bowtie2. Window widths of 1500 bp was used for Cas9 ChIP-seq, and 2500 bp was used for MRE11 ChIP-seq.

### Nucleotide composition analysis of region surrounding gRNA on-target sites

The local genomic sequences for each expected on-target site for ‘CT’, ‘GG’, and ‘TA’ gRNAs were obtained, then aligned by the Cas9 cut site with PAM always oriented downstream of the cut). At each base pair position relative to the cut site, the nucleotide identity (A, C, G, T) was tallied and/or displayed. This analysis was performed +/- 500 bp from the cut sites.

### Calculating enrichment for MRE11, Cas9, γH2AX, and 53BP1 ChIP-seq

Custom code using the pysam package determines the number of reads, per million of reads, in specific window widths centered at all cut sites. A 200 kb window was used for both 53BP1 and γH2AX. A 2500 bp window was used for MRE11, and 1500 window was used for Cas9. For MRE11 and Cas9, additional code analyzes the exact read positions and determines if a paired-end sequencing read fragment spans the cut site (‘span’), or if a sequenced DNA fragment begins within 5 bp from the cut site (‘abut’). To determine ‘dist+4’, ‘dist-4’, ‘prox+4’, or ‘prox-4’, the code further analyzes the DNA fragment position according to the rules specified for these read species.

### Enrichment profiles for MRE11 and Cas9 ChIP-seq (also spanning ATAC-seq) at base-pair resolution

At each genomic position in a window centered at each cut site, each paired-end read within this window is retrieved. The number of paired-end reads that map to each base pair is tallied. The middle region of PE read fragment that is not likely to be sequenced is also included in this tally. A 2500 bp window was used for MRE11 and 1500 window was used for Cas9.

We used the same protocol to evaluate the profiles of ATAC-seq reads that span the cut site, using a 3 kb window.

### Enrichment profiles for γH2AX, 53BP1, and ATAC-seq at window widths

To obtain profiles of γH2AX and 53BP1, calculate the number of sequencing reads (per million total sequencing reads) in each 10 kb window from the cut site, extending to 2 mb both upstream and downstream of cut sites.

For ATAC-seq, calculate reads per million in a 4 bp sliding window incremented every 1 bp, extending to 1.5 kb both upstream and downstream of cut sites.

To determine wider levels of potential ATAC-seq enrichment, we used the same function to calculate reads per million in each 1 kb window from the cut site, extending to 50 kb both upstream and downstream of cut sites.

### Genome-wide Cas9 binding from dCas9 ChIP-seq

Used macs2 to find all dCas9 binding peaks, using a no-Cas9 sample for negative control. ‘-f BAMPE’ indicates paired-end format, ‘-g hs’ species the genome size of homo sapiens (2.7e9 bp).

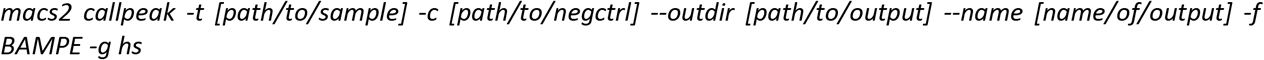

Next, for each macs2 discovered peak with fold enrichment >= 4, a custom algorithm attempts to identify the target sequence position for Cas9 binding or cleavage that best explains the peak. This may be problematic for target sites with multiple mismatches. We use the following assumption to simplify the problem: (1) there is only one correct Cas9 binding/cleavage sequence within the 400 bp window of the macs2-predicted peak center, and (2) the correct Cas9 binding/cleavage sequence is one with the fewest mismatches.

### Enrichment measurements of epigenetic markers

We use the following publicly available epigenetic datasets, all from HEK293 cell lines: ATAC-seq (SRR6418075), DNaseI (ENCFF120XFB), H3K4me1 (ENCFF909ESY), H3K4me3 (ENCFF912BYL), H3K9me3 (ENCFF141ZEQ), H3K27ac (ENCFF588KSR), H3K36me3 (ENCFF593SUW), MNase-seq (ERR2403161), and RNA-seq (SRR5627161). Datasets starting with ENCFF can be found and downloaded from ENCODE (https://www.encodeproject.org/). Datasets starting with SRR or ERR can be found and downloaded from NIH’s SRA (https://www.ncbi.nlm.nih.gov/sra).

For enrichment, we use a 50 kb radius for RNA-seq, H3K4me1, H3K4me3, H3K9me3, H3K27ac, and H3K36me3, a 50 bp radius for DNaseI and ATAC-seq, and a 10 bp radius for MNase-seq. The number of reads that are found in each specified window width is outputted, normalized by the total sequencing reads per million.

### Machine learning model

We used the Random Forest Regressor from scikit-learn^57^. For mismatch information, features were obtained from one-hot encoding of mismatch state at each position along the protospacer. For epigenetic information, the reads per million enrichment was directly used as features. The predicted output is the level of dCas9 binding or MRE11 enrichment, also measured as reads per million. The machine learning model was trained using 5-fold cross validation on a training dataset comprised of a random 70% of the total dataset. The remaining 30% was used for evaluation and featured in these figures comparing predicted vs actual values.

### ATAC-seq sequencing read length distributions

For each paired-end ATAC-seq read fragment in a 3 kb window centered at all Cas9 on-target sites, its length was recorded. The distribution of DNA length across all target sites, along with exponential decay curve fitting, was computed on Microsoft Excel.

