## Supplementary Material for "Massively parallel genomic perturbations with multi-target CRISPR reveal new insights on Cas9 activity and DNA damage responses at endogenous sites"

**Supplementary Figures 1 to 6**

**Supplementary Tables 1 to 8**

**Supplementary Data 1 to 15**

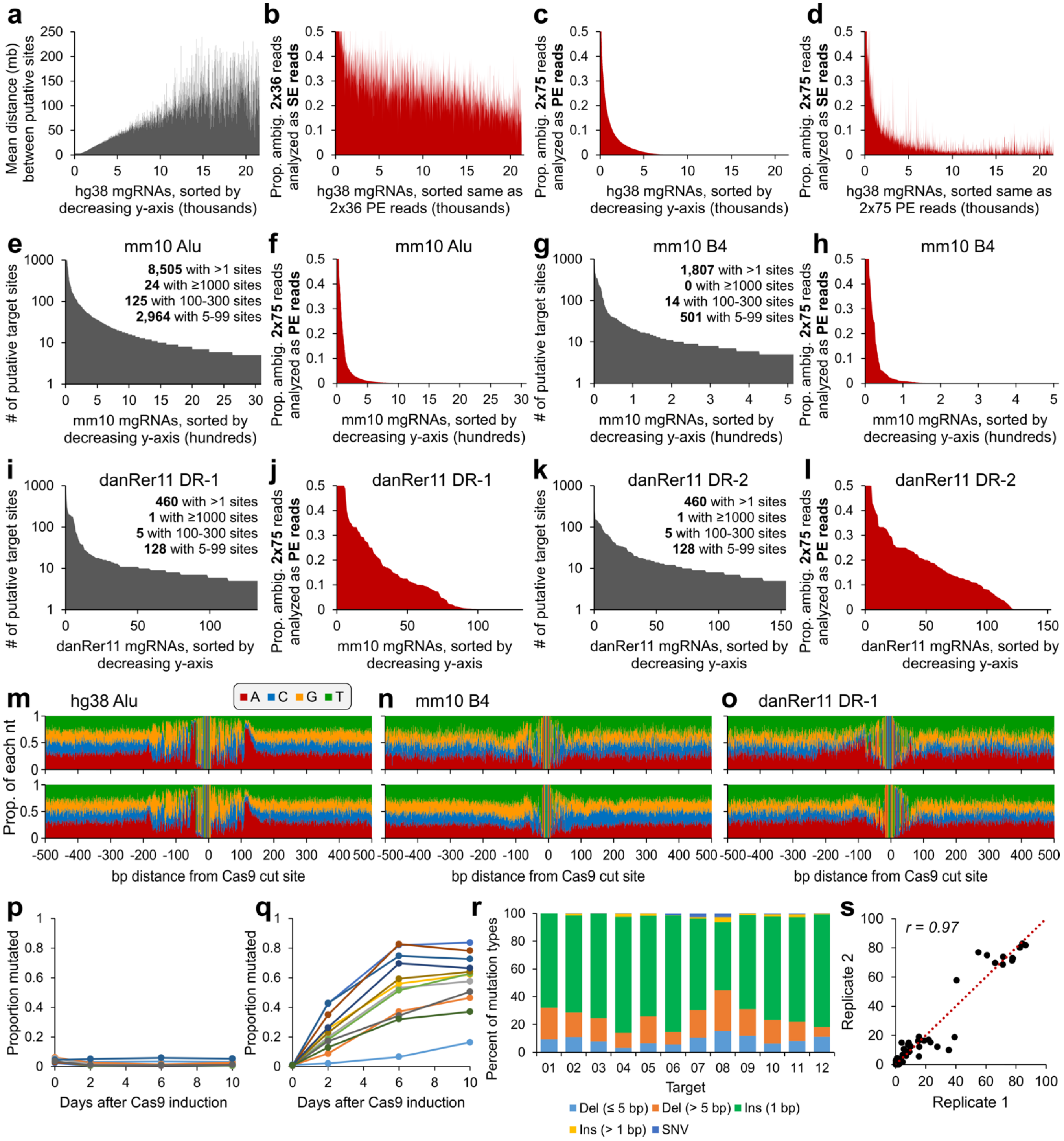
a
Mean distance (mb) between putative sites
hg38 mgRNAs, sorted by decreasing y-axis (thousands)
b
Prop. ambig. 2x36 reads analyzed as SE reads
hg38 mgRNAs, sorted same as 2x36 PE reads (thousands)
c
Prop. ambig. 2x75 reads analyzed as PE reads
hg38 mgRNAs, sorted by decreasing y-axis (thousands)
d
Prop. ambig. 2x75 reads analyzed as SE reads
hg38 mgRNAs, sorted same as 2x75 PE reads (thousands)
e
mm10 Alu
8,505 with >1 sites
24 with ≥1000 sites
125 with 100-300 sites
2,964 with 5-99 sites
### of putative target sites
mm10 mgRNAs, sorted by decreasing y-axis (hundreds)
f
mm10 Alu
Prop. ambig. 2x75 reads analyzed as PE reads
mm10 mgRNAs, sorted by decreasing y-axis (hundreds)
g
mm10 B4
1,807 with >1 sites
0 with ≥1000 sites
14 with 100-300 sites
501 with 5-99 sites
### of putative target sites
mm10 mgRNAs, sorted by decreasing y-axis (hundreds)
h
mm10 B4
Prop. ambig. 2x75 reads analyzed as PE reads
mm10 mgRNAs, sorted by decreasing y-axis (hundreds)
i
danRer11 DR-1
460 with >1 sites
1 with ≥1000 sites
5 with 100-300 sites
128 with 5-99 sites
### of putative target sites
danRer11 mgRNAs, sorted by decreasing y-axis
j
danRer11 DR-1
Prop. ambig. 2x75 reads analyzed as PE reads
mm10 mgRNAs, sorted by decreasing y-axis
k
danRer11 DR-2
460 with >1 sites
1 with ≥1000 sites
5 with 100-300 sites
128 with 5-99 sites
### of putative target sites
danRer11 mgRNAs, sorted by decreasing y-axis
l
danRer11 DR-2
Prop. ambig. 2x75 reads analyzed as PE reads
danRer11 mgRNAs, sorted by decreasing y-axis
m
hg38 Alu
A C G T
n
mm10 B4
o
danRer11 DR-1
Prop. of each nt
bp distance from Cas9 cut site
p
Proportion mutated
Days after Cas9 induction
q
Proportion mutated
Days after Cas9 induction
r
Percent of mutation types
Target
01 02 03 04 05 06 07 08 09 10 11 12
Del (≤ 5 bp) Del (> 5 bp) Ins (1 bp)
Ins (> 1 bp) SNV
s
r = 0.97
Replicate 2
Replicate 1

**Figure S1: *in silico* characterization of multi-target gRNAs**

**a,** For the unique target sequences shown in Fig. 1b, we plotted the mean distance (in megabases) between adjacent putative on-target sites on the same chromosome. The order of sequences along the x-axis is the same as in Fig. 1b.

**b,** Same as Fig. 1e, but interpreted as two single end (SE) 36bp ChIP-seq reads.

**c,** Same as Fig. 1e, but from simulated PE 2x75bp ChIP-seq reads. Target sequences (x-axis) are sorted by decreasing proportion of ambiguous reads (y-axis).

**d,** Same as Fig. S1c, but interpreted as two single end (SE) 75bp ChIP-seq reads.

**e-f,** Same as Fig. 1b and Fig. 1e, respectively, but for the Alu SINE in the mouse mm10 genome.

**g-h,** Same as Fig. 1b and Fig. 1e, respectively, but for the B4 SINE in the mouse mm10 genome.

**i-j,** Same as Fig. 1b and Fig. 1e, respectively, but for the DR-1 SINE in the zebrafish danRer11 genome.

**k-l,** Same as Fig. 1b and Fig. 1e, respectively, but for the DR-2 SINE in the zebrafish danRer11 genome.

**m-o,** Same as Fig. 1g, but for two different mgRNAs from the (m) Alu SINE in the human hg38 genome, (n) B4 SINE in the mouse mm10 genome, and (o) DR-1 SINE in the zebrafish danRer11 genome.

**p,** No-dox control mutation curve. Dox-inducible Cas9 cells genomically integrated with a 10-target mgRNA (same cells as in Fig. 2a) were grown and passaged without dox exposure. Cells were harvested at different time points (0, 2, 6 and 10 days). Figure details are the same as in Fig. 2a.

**q,** Mutation curve of a 20-target mgRNA in HeLa cells with dox-inducible Cas9. Figure details are the same as in Fig. 2a and Fig. S1p.

**r,** Mutation signatures of a 20-target mgRNA, outcomes determined for 10-day samples from Fig. S1q. Figure details are the same as in Fig. 2b.

**s,** Same reproducibility analysis and figure details as in Fig. 2c but using the 20-target mgRNA data.

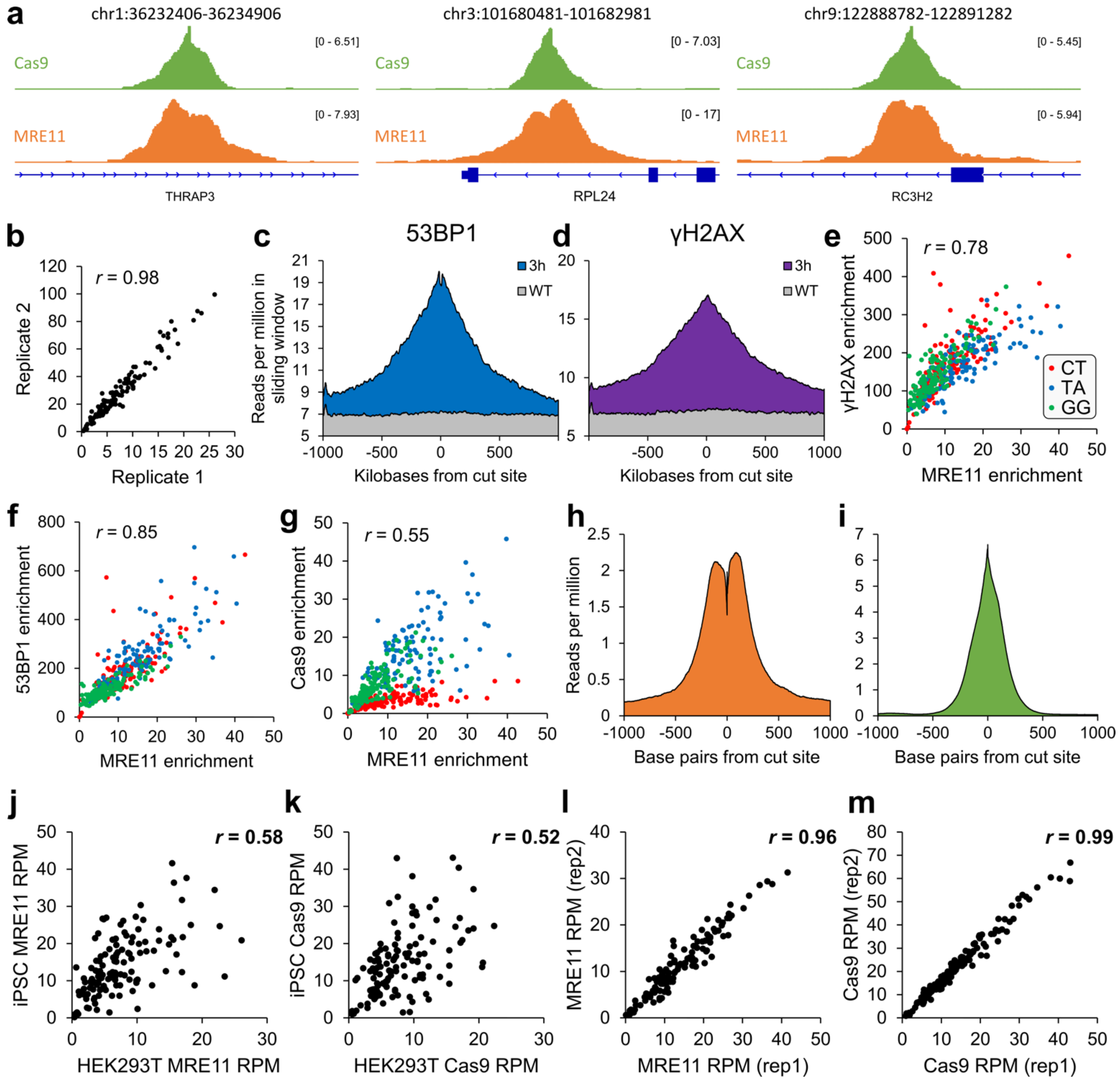

a
chr1:36232406-36234906
chr3:101680481-101682981
chr9:122888782-122891282
Cas9
MRE11
[0 - 6.51]
[0 - 7.93]
[0 - 7.03]
[0 - 17]
[0 - 5.45]
[0 - 5.94]
THRAP3
RPL24
RC3H2
b
r = 0.98
Replicate 2
Replicate 1
c
53BP1
3h
WT
Reads per million in sliding window
Kilobases from cut site
d
γH2AX
e
r = 0.78
γH2AX enrichment
MRE11 enrichment
CT
TA
GG
f
r = 0.85
53BP1 enrichment
g
r = 0.55
Cas9 enrichment
h
Reads per million
Base pairs from cut site
i
j
r = 0.58
iPSC MRE11 RPM
HEK293T MRE11 RPM
k
r = 0.52
iPSC Cas9 RPM
HEK293T Cas9 RPM
l
r = 0.96
MRE11 RPM (rep2)
MRE11 RPM (rep1)
m
r = 0.99
Cas9 RPM (rep2)
Cas9 RPM (rep1)

**Figure S2: Enrichment quantification of Cas9 and DNA repair factor ChIP-seq at on-target sites**

**a,** Coordinates of cut sites in hg38 are listed in the title of each panel. Numerical value of enrichment is reported as reads per million, *i.e.* at each genomic coordinate, the number of paired-end ChIP-seq read fragments that span this coordinate, per million aligned ChIP-seq reads.

**b,** Correlation between two biological replicates of MRE11 ChIP-seq. Both axes have units of reads per million (RPM) enrichment in a 2.5 kb window centered at each cut site.

**c-d,** Average profiles of (c) 53BP1 and (d) γH2AX enrichment in a 2 megabase window centered at the cut sites.

**e-g,** Plots of relationships between (e) MRE11 and γH2AX, (f) MRE11 and 53BP1, and (g) MRE11 and Cas9 at each target site of the three 3 different target sequences (‘CT’ – red, ‘TA’ – blue, ‘GG’ – green).

**h-i,** Average profile of (h) MRE11 and (i) Cas9 enrichment in a 2000 bp window centered at all cut sites in WTC-11 induced pluripotent stem cells (iPSCs).

**j-k,** Plot of (j) MRE11 and (k) Cas9 ChIP-seq enrichment around all target sites, between HEK293T cells and iPSCs.

**l-m,** Plot of (l) MRE11 and (m) Cas9 ChIP-seq enrichment around all target sites, between iPSC biological replicates.

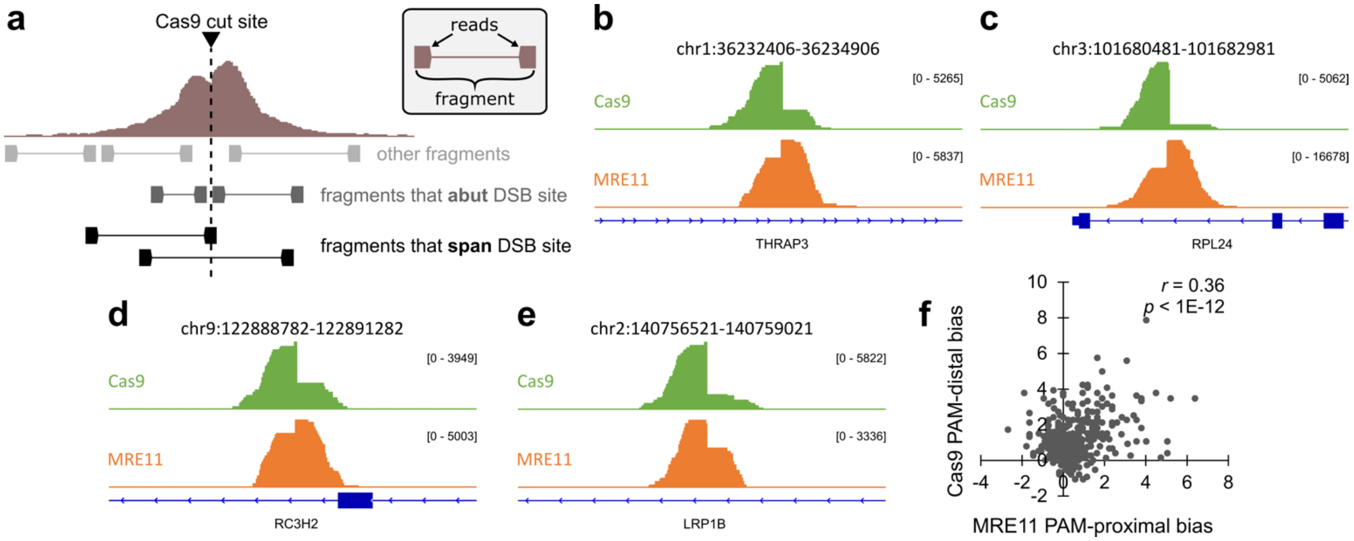


**Figure S3: Fragment and mismatch analysis for MRE11 and Cas9 ChIP-seq**

**a,** Conceptual schematic of ChIP-seq analysis for paired-end reads, distinguishing between read fragments that abut or span the DSB site. Reads abut the DSB site if within 5 bp from the cut site.

**b-e,** For all 4 target sites, the PAM-distal side is oriented on the left. (b-d) The first three examples exhibit Cas9 PAM-distal bias and MRE11 PAM-proximal bias. However, the last example (e) in chr2:140756521-140759021 illustrates an exception where both Cas9 and MRE11 exhibit PAM-distal bias.

**f,** For each target site, plotted the MRE11 PAM-proximal bias versus Cas9 PAM-distal bias, observing that the two quantities are correlated. PAM-proximal bias is defined as reads per million (RPM) on the PAM-proximal side minus RPM on the PAM-distal side, and vice versa for PAM-distal bias.

**a**

| | | 00 | 01 (d) | 01 (p) | 02 (d) | 02 (m) | 02 (p) | 03 | 04 | 05 | 06 | 07 | 08 | 09 | 10 | 11 | 12 |
|---|---|---|---|---|---|---|---|---|---|---|---|---|---|---|---|---|---|
| MRE11 | rep 1 | 27.60 | 17.33 | 1.41 | 7.59 | 1.43 | 0.83 | 0.99 | 0.92 | 0.70 | 0.79 | 0.90 | 0.82 | 0.85 | 0.93 | 0.89 | 0.46 |
| | rep 2 | 20.63 | 12.84 | 1.27 | 6.55 | 1.31 | 0.92 | 1.11 | 1.10 | 1.22 | 1.14 | 1.04 | 1.05 | 1.00 | 1.19 | 1.19 | 1.13 |
| dCas9 | rep 1 | 282.03 | 100.45 | 16.91 | 66.98 | 8.86 | 12.42 | 11.26 | 23.17 | 30.05 | 28.32 | 20.25 | 18.30 | 14.01 | 10.67 | 5.93 | 8.64 |
| | rep 2 | 294.27 | 99.25 | 17.47 | 62.90 | 7.78 | 11.54 | 9.22 | 17.48 | 29.71 | 25.26 | 17.76 | 16.58 | 12.74 | 9.43 | 5.04 | 8.88 |

**b**

| | H3K4me1 | H3K4me3 | H3K9me3 | H3K27ac | H3K36me3 | DNase I | MNase | ATAC-seq | RNA-seq |
|---|---|---|---|---|---|---|---|---|---|
| H3K4me1 | 1.00 | | | | | | | | |
| H3K4me3 | 0.49 | 1.00 | | | | | | | |
| H3K9me3 | 0.20 | -0.14 | 1.00 | | | | | | |
| H3K27ac | 0.73 | 0.84 | -0.04 | 1.00 | | | | | |
| H3K36me3 | 0.32 | 0.64 | -0.17 | 0.56 | 1.00 | | | | |
| DNase I | 0.15 | 0.05 | -0.10 | 0.11 | 0.00 | 1.00 | | | |
| MNase | -0.02 | -0.04 | -0.20 | -0.06 | -0.04 | 0.13 | 1.00 | | |
| ATAC-seq | 0.30 | 0.09 | 0.05 | 0.28 | 0.11 | 0.40 | 0.01 | 1.00 | |
| RNA-seq | 0.39 | 0.71 | -0.18 | 0.67 | 0.74 | 0.00 | 0.03 | 0.12 | 1.00 |

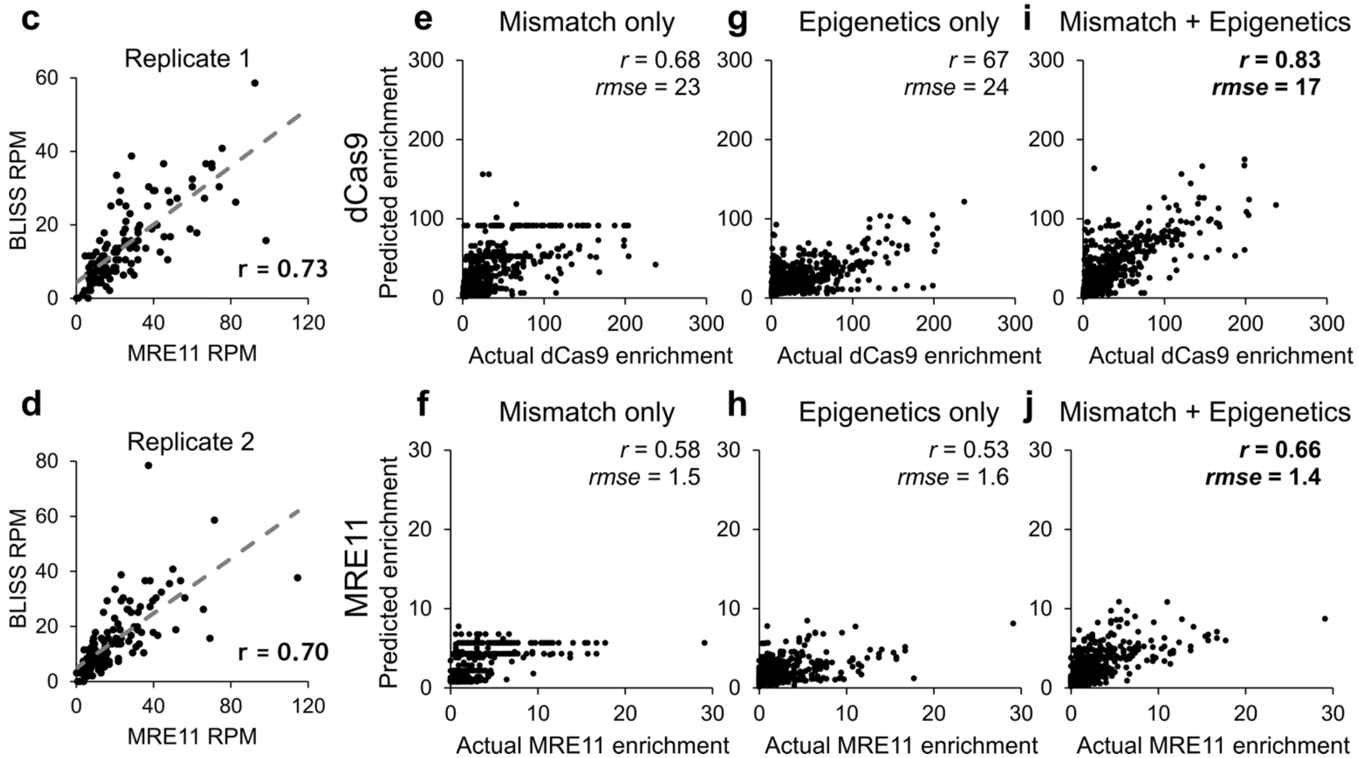


**Figure S4: Further correlation analysis and machine learning modeling at 30 min**

**a,** Mean quantification of Fig. 4b-c for two biological replicates.

**b,** Correlation coefficients between enrichment of public epigenetic datasets at all on-target sites targeted by the 'GG' multi-target gRNA.

**c-d,** Two replicates of correlation between MRE11 and BLISS enrichment in 2500 bp windows centered around all cut sites.

**e-j,** Same as Fig. 4f-k but for samples evaluated at 30 minutes after (d)Cas9 delivery.

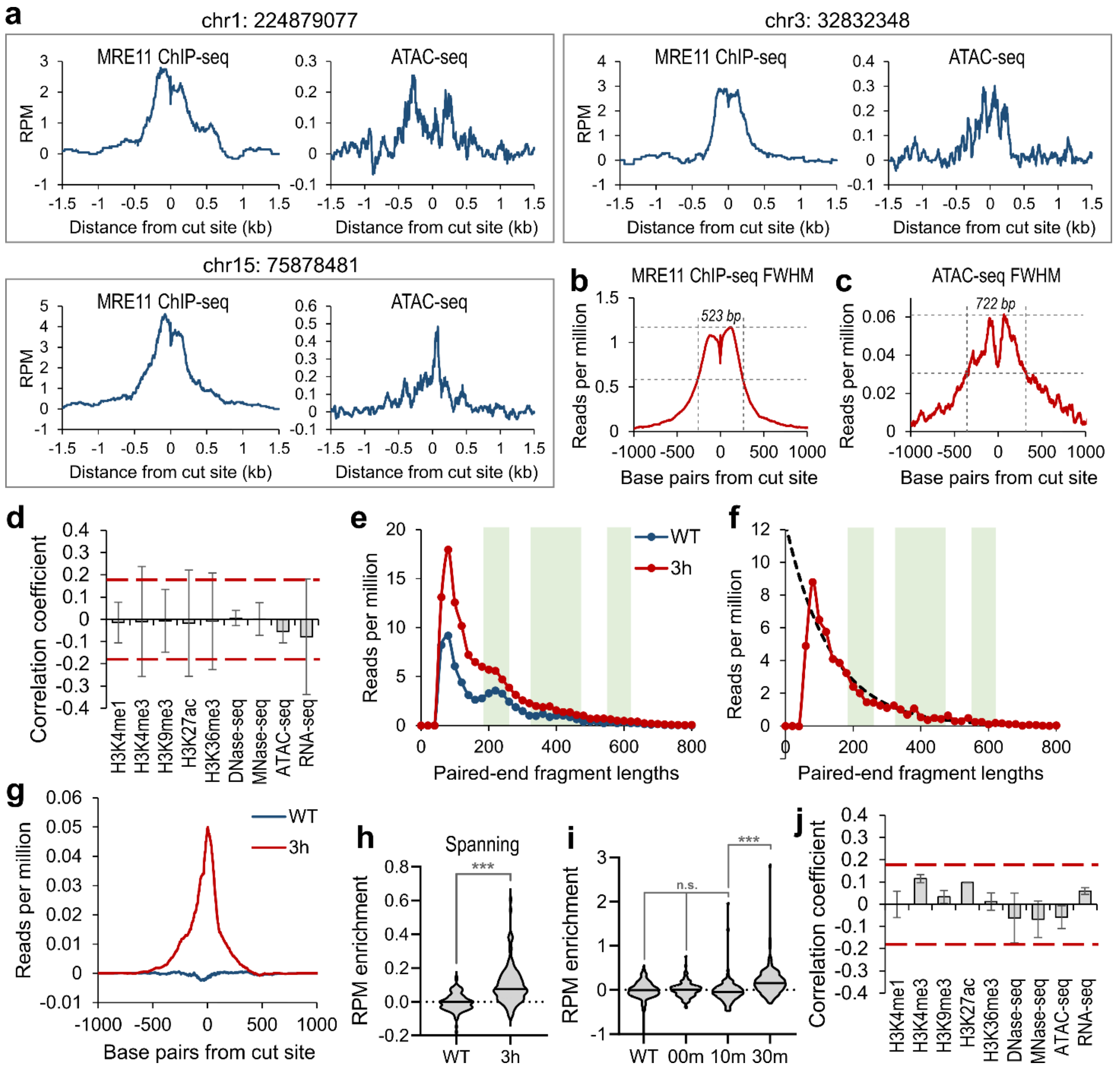


**Figure S5: Local ATAC-seq enrichment after Cas9 DNA damage**

**a,** Three representative cut sites are labeled from the human genome hg38. The MRE11 enrichment from MRE11 ChIP-seq and background-subtracted chromatin accessibility increase from ATAC-seq are both plotted.

**b-c,** Measurement of full width at half maximum (FWHM) for MRE11 ChIP-seq enrichment (523 bp) versus MRE11 enrichment (722 bp) averaged across all on-target cut sites.

**d,** Correlation between 9 epigenetic markers and MRE11-normalized excess ATAC-seq enrichment due to Cas9-mediated DNA damage.

**e-h,** Second biological replicate of Fig. 5d-g.

**i,** Violin plots of background-subtracted ATAC-seq reads per million (RPM) enrichment at all on-target sites. ‘WT’ indicates Cas9-negative cells. ‘00m’, ‘10m’, and ‘30m’ indicate cells with vfCRISPR Cas9/cgRNA without light exposure, 10 minutes after light exposure, and 30 minutes after light exposure, respectively.

**j,** Proportional change in background-subtracted MRE11 RPM enrichment at all on-target sites is not correlated with any evaluated epigenetic markers.

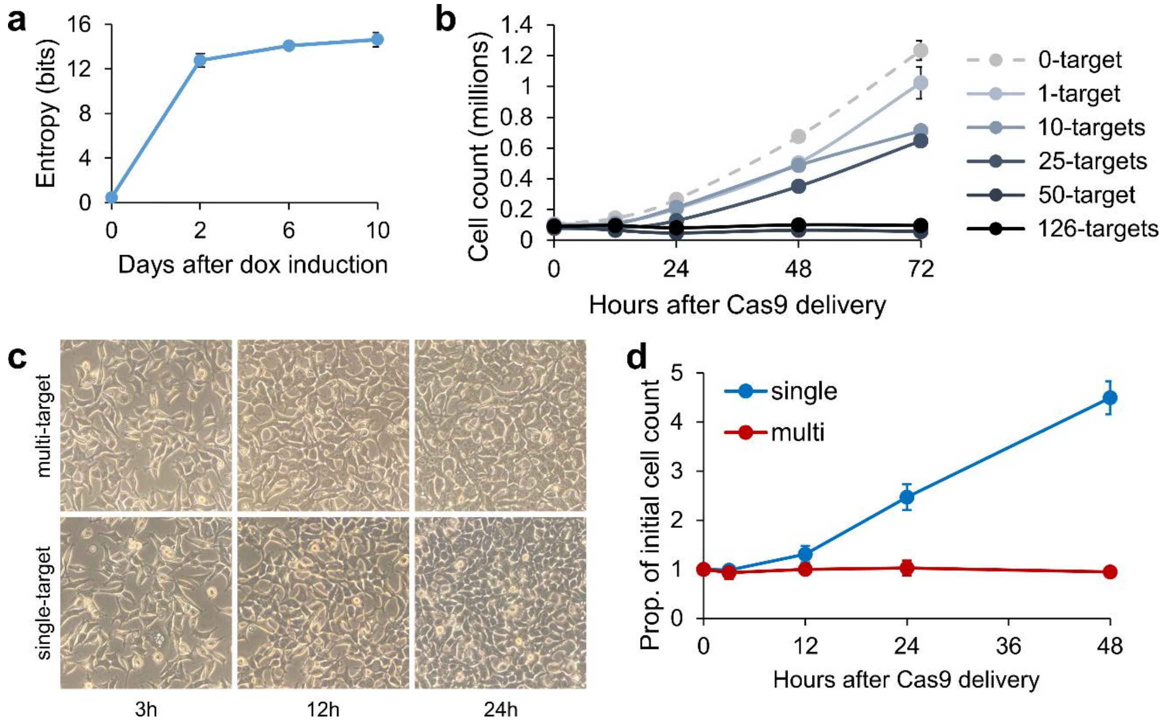


**Figure S6: Cellular barcoding and quantification of cell growth with multi-target CRISPR**

**a,** Cumulative Shannon entropy for the 10-target mgRNAs across the 8 on-target sites with detectable indels. Higher entropy corresponds to higher mutation diversity that is necessary for effective cellular barcoding.

**b,** Cell counts over time (12h, 24h, 48h, 72h) after delivery of Cas9 into HEK293T cells with mgRNA targeting various numbers of expected genome-wide sites (0, 1, 10, 25, 50, 126 expected sites). Data points are the average of two biological replicates and error bars are the standard deviation.

**c,** Representative microscopy images of HEK293T cells at different time points (3h, 12h, 24h) after electroporation of Cas9 with multi-target (‘GG’) versus single-target (targeting *ACTB*) gRNA.

**d,** Quantification of Fig. S6c.

**Table S1: Our study using mgRNAs versus other methods that characterize Cas9 activity**

| | model system | | | effect type | | measured aspect of genome editing | | | |
|---|---|---|---|---|---|---|---|---|---|
| | DNA only | transformed cells | primary cells | sequence | epigenetics | binding | cleavage | damage response | outcome (indels) |
| this paper (mgRNAs) | - | + | + | + | +++ | + | + | + | + |
| Lazzarotto et al., 2020 (CHANGE-seq) | + | - | - | +++ | ++ | - | + | - | - |
| Boyle et al., 2021 | + | - | - | +++ | - | + | + | - | - |
| Gisler et al., 2019 | - | + | - | +++ | + | - | - | - | + |
| Chakrabarti et al., 2019 | - | + | - | +++ | + | - | - | - | + |
| Shen et al., 2018 (inDelphi) | - | + | - | +++ | - | - | - | - | + |
| Allen et al., 2018 | - | + | - | +++ | - | - | - | - | + |
| van Overbeek et al., 2016 | - | + | - | +++ | - | - | - | - | + |

**Supplementary Table 2: crRNA and tracrRNA sequences**

- tracrRNA was purchased as Alt-R® CRISPR-Cas9 tracrRNA from Integrated DNA Technologies (IDT)
- IDT_* crRNAs were purchased as 36 nt Alt-R® CRISPR-Cas9 crRNAs from IDT
- cg_* crRNAs are crRNAs for vfCRISPR (*Liu et al., 2020*), purchased as a custom order from BioSynthesis
- pc_* crRNAs are crRNAs for rapid Cas9 deactivation (*Zou et al., 2021*), purchased as a custom order from IDT
- The red bases hybridize with target DNA

| **Name** | **Sequence (5′ to 3′)** |
|---|---|
| tracrRNA | AGCAUAGCAAGUUAAAAUAAGGCUAGUCCGUUAUCAACUUGAAAAAGUGGCACCGAGUCGGUGCUUU |
| IDT_AluGG | **CCT GTA GTC CCA GCT ACT GG**G UUU UAG AGC UAU GCU |
| IDT_AluCT | **CCT GTA GTC CCA GCT ACT CT**G UUU UAG AGC UAU GCU |
| IDT_AluTA | **CCT GTA GTC CCA GCT ACT TA**G UUU UAG AGC UAU GCU |
| cg_AluGG | **CC[NPOM-dT] G[NPOM-dT]A G[NPOM-dT]C CCA GCU ACU GG**G UUU UAG AGC UAU GCU GUU UUG |
| Pc_AluGG | **CCU GU[PC-Linker] GUC CCA GCU ACU GG**G UUU UAG AGC UAU GCU GUU UUG |

**Supplementary Table 3: Oligos for preparing ChIP sequencing libraries**

| **Name** | **Sequence (5′ to 3′) \| Red marks the i7 6 bp index sequence** |
|---|---|
| MNase_F | /5Phos/GATCGGAAGAGCACACGTCT |
| MNase_R | ACACTCTTTCCCTACACGACGCTCTTCCGATC*T |
| PE_i5 | AATGATACGGCGACCACCGAGATCTACACTCTTTCCCTACACGACGCTCTTCCGATC*T |
| PE_i701 | CAAGCAGAAGACGGCATACGAGAT**CGTGAT**GTGACTGGAGTTCAGACGTGTGCTCTTCCGATC*T |
| PE_i702 | CAAGCAGAAGACGGCATACGAGAT**ACATCG**GTGACTGGAGTTCAGACGTGTGCTCTTCCGATC*T |
| PE_i703 | CAAGCAGAAGACGGCATACGAGAT**GCCTAA**GTGACTGGAGTTCAGACGTGTGCTCTTCCGATC*T |
| PE_i704 | CAAGCAGAAGACGGCATACGAGAT**TGGTCA**GTGACTGGAGTTCAGACGTGTGCTCTTCCGATC*T |
| PE_i705 | CAAGCAGAAGACGGCATACGAGAT**CACTGT**GTGACTGGAGTTCAGACGTGTGCTCTTCCGATC*T |
| PE_i706 | CAAGCAGAAGACGGCATACGAGAT**ATTGGC**GTGACTGGAGTTCAGACGTGTGCTCTTCCGATC*T |
| PE_i707 | CAAGCAGAAGACGGCATACGAGAT**GATCTG**GTGACTGGAGTTCAGACGTGTGCTCTTCCGATC*T |
| PE_i708 | CAAGCAGAAGACGGCATACGAGAT**TCAAGT**GTGACTGGAGTTCAGACGTGTGCTCTTCCGATC*T |
| PE_i709 | CAAGCAGAAGACGGCATACGAGAT**CTGATC**GTGACTGGAGTTCAGACGTGTGCTCTTCCGATC*T |
| PE_i710 | CAAGCAGAAGACGGCATACGAGAT**AAGCTA**GTGACTGGAGTTCAGACGTGTGCTCTTCCGATC*T |
| PE_i711 | CAAGCAGAAGACGGCATACGAGAT**GTAGCC**GTGACTGGAGTTCAGACGTGTGCTCTTCCGATC*T |
| PE_i712 | CAAGCAGAAGACGGCATACGAGAT**TACAAG**GTGACTGGAGTTCAGACGTGTGCTCTTCCGATC*T |
| PE_i713 | CAAGCAGAAGACGGCATACGAGAT**ATCAGT**GTGACTGGAGTTCAGACGTGTGCTCTTCCGATC*T |
| PE_i714 | CAAGCAGAAGACGGCATACGAGAT**AGGAAT**GTGACTGGAGTTCAGACGTGTGCTCTTCCGATC*T |
| PE_i715 | CAAGCAGAAGACGGCATACGAGAT**ATTCCG**GTGACTGGAGTTCAGACGTGTGCTCTTCCGATC*T |
| PE_i716 | CAAGCAGAAGACGGCATACGAGAT**CCACTC**GTGACTGGAGTTCAGACGTGTGCTCTTCCGATC*T |

| | |
|---|---|
| PE_i717 | CAAGCAGAAGACGGCATACGAGAT**CGATTA**GTGACTGGAGTTCAGACGTGTGCTCTTCCGATC*T |
| PE_i718 | CAAGCAGAAGACGGCATACGAGAT**CTTCGA**GTGACTGGAGTTCAGACGTGTGCTCTTCCGATC*T |
| PE_i719 | CAAGCAGAAGACGGCATACGAGAT**GAATGA**GTGACTGGAGTTCAGACGTGTGCTCTTCCGATC*T |
| PE_i720 | CAAGCAGAAGACGGCATACGAGAT**GCGGAC**GTGACTGGAGTTCAGACGTGTGCTCTTCCGATC*T |
| PE_i721 | CAAGCAGAAGACGGCATACGAGAT**GGAACT**GTGACTGGAGTTCAGACGTGTGCTCTTCCGATC*T |
| PE_i722 | CAAGCAGAAGACGGCATACGAGAT**TAGTTG**GTGACTGGAGTTCAGACGTGTGCTCTTCCGATC*T |
| PE_i723 | CAAGCAGAAGACGGCATACGAGAT**TCGGGA**GTGACTGGAGTTCAGACGTGTGCTCTTCCGATC*T |
| PE_i724 | CAAGCAGAAGACGGCATACGAGAT**TCTGAG**GTGACTGGAGTTCAGACGTGTGCTCTTCCGATC*T |

#### Supplementary Table 4: Oligos for BLISS

| Name | Sequence (5' to 3') |
|---|---|
| MNase_F | /5Phos/GATCGGAAGAGCACACGTCT |
| MNase_R | ACACTCTTTCCCTACACGACGCTCTTCCGATC*T |

#### Supplementary Table 5: Oligos for cloning mgRNAs into lentiviral vector.

| Name | Sequence (5' to 3') |
|---|---|
| Ct10_mgRNA_F | CACCGCCAGGCTGGAGTGCAGTGCT |
| Ct10_mgRNA_R | AAACAGCACTGCACTCCAGCCTGGC |
| Ct20_mgRNA_F | CACCGGCACTCCAGCCTGGGTTACA |
| Ct20_mgRNA_R | AAACTGTAACCCAGGCTGGAGTGCC |

#### Supplementary Table 6: Oligos for sequencing mgRNA targets. PCR-1

| **10-target mgRNA** | |
|---|---|
| **Name** | **Sequence (5' to 3')** |
| ct10_T1_Out_F | AAGGGAGGCTGGGAAATGTG |
| ct10_T1_Out_R | GCATCCCTCAGCCTCAGTTT |
| ct10_T2_Out_F | TCCTGGATCTGGCCTAAGCT |
| ct10_T2_Out_R | GGGCAGCATAGCAAGATCCT |
| ct10_T3_Out_F | TGAAGCAGCTGCAGAAGAACT |
| ct10_T3_Out_R | GTGGTGGCTCACGTCTGTAA |
| ct10_T4_Out_F | TGGTTTTGGAGCGCTCTTCA |
| ct10_T4_Out_R | CCCGAAGAGCAAAGACAGGT |
| ct10_T5_Out_F | TCATGGGGTCACCTTATACCA |
| ct10_T5_Out_R | TTACTCTGGCTTGCAGGGTG |
| ct10_T6_Out_F | AAACAGACACAAAAAGCAGTCA |
| ct10_T6_Out_R | GCCAAGGCAGGAGAGTCATT |
| ct10_T7_Out_F | GGTCTCAAACTCCTCGGCTC |
| ct10_T7_Out_R | CATGGTGACTTGTCCCAGCA |
| ct10_T8_Out_F | TGGACTTCTCAGCCTCCAGA |
| ct10_T8_Out_R | GCATCCCCAAGTGGTTCTGA |
| ct10_T9_Out_F | GTGGGAAGGATGAGGCAGAG |
| ct10_T9_Out_R | TTGAAGGGGTTTCTGGAGGC |
| **20-target mgRNA** | |
| ct20_T1_Out_F | TGGATCACGAGGTCAGGAGA |
| ct20_T1_Out_F | AGTGGGGGTGGTTAGAGAGA |
| ct20_T2_Out_F | CAGCCTGGGCAACTTAGTGA |
| ct20_T2_Out_R | TGGTGGGTTTTGCTTTCCCT |
| ct20_T3_Out_F | TAGCATTTTGGGAGGCCGAG |
| ct20_T3_Out_R | ACACACACTGAAAAACGCAA |
| ct20_T4_Out_F | GGGCAGATCATGAGGTCGAG |
| ct20_T4_Out_R | ATACGTAGGAGGCATGCTCC |
| ct20_T5_Out_F | TTGAAACAAGCCTGGCCAAC |

| | |
|---|---|
| ct20_T5_Out_R | GAGCAAGGGACAGAGCAGTT |
| ct20_T6_Out_F | GTGGGCAGATCACAAGGTCA |
| ct20_T6_Out_R | TGGCAAATAAGAGAATGTGGGG |
| ct20_T7_Out_F | GGAGTTCAAGACCAGTCCGG |
| ct20_T7_Out_R | GAACCTCCTGTGTCCCTGTG |
| ct20_T8_Out_F | GACCAGCTTGGCCAAAATGG |
| ct20_T8_Out_R | TGGTGCTAGTTATACCAGCCC |
| ct20_T9_Out_F | ATCCCAGCACTTTGAGAGGC |
| ct20_T9_Out_R | ACCCAGATAGAGCCCGTCTT |
| ct20_T10_Out_F | GATCGCTTGAGCTCAGGTGT |
| ct20_T10_Out_R | CCCCAGAATTCGACTGTCAGG |
| ct20_T11_Out_F | AAGTTCAAGACCAGCCTGGG |
| ct20_T11_Out_R | GAGACGGGGTTTCACCATGT |
| ct20_T12_Out_F | CAGCCTGGGCAACATAGTGA |
| ct20_T12_Out_R | TTCCCACCTAAACCTTGCCC |

#### Supplementary Table 7: Oligos for sequencing mgRNA targets. PCR-2

| **Name** | **Sequence (5′ to 3′)** |
|---|---|
| **10-target mgRNA** | |
| ct10_T1_In_F | TCGTCGGCAGCGTCAGATGTGTATAAGAGACAGGTCAGGAGTTCGAGACCAGC |
| ct10_T1_In_R | GTCTCGTGGGCTCGGAGATGTGTATAAGAGACAGAGGGCAGTGGTAAGGAGGAT |
| ct10_T2_In_F | TCGTCGGCAGCGTCAGATGTGTATAAGAGACAGACAGCTTGGGGTTACACAGG |
| ct10_T2_In_R | GTCTCGTGGGCTCGGAGATGTGTATAAGAGACAGGCCTCACAATGGCCCTATGA |
| ct10_T3_In_F | TCGTCGGCAGCGTCAGATGTGTATAAGAGACAGGCAGAGGCATGTGGATCACT |
| ct10_T3_In_R | GTCTCGTGGGCTCGGAGATGTGTATAAGAGACAGGCCAGTGGACTTTAGGCAAA |
| ct10_T4_In_F | TCGTCGGCAGCGTCAGATGTGTATAAGAGACAGAGCCAGTGTGTGACTTAGGC |
| ct10_T4_In_R | GTCTCGTGGGCTCGGAGATGTGTATAAGAGACAGGAGATCAGCCTGGGCAACAT |
| ct10_T5_In_F | TCGTCGGCAGCGTCAGATGTGTATAAGAGACAGCGCTGAGGCAGGTAGATCAC |
| ct10_T5_In_R | GTCTCGTGGGCTCGGAGATGTGTATAAGAGACAGTGCCATCACTGGGTCTAATGG |
| ct10_T6_In_F | TCGTCGGCAGCGTCAGATGTGTATAAGAGACAGATTAGCTGGGTGTGGTGACG |
| ct10_T6_In_R | GTCTCGTGGGCTCGGAGATGTGTATAAGAGACAGTCAGCTGTTTGATCTCAGGCA |
| ct10_T7_In_F | TCGTCGGCAGCGTCAGATGTGTATAAGAGACAGAGAGCCTGGACAAAATGGTGA |
| ct10_T7_In_R | GTCTCGTGGGCTCGGAGATGTGTATAAGAGACAGGGGTCTAGCAAGTGGGCAAT |
| ct10_T8_In_F | TCGTCGGCAGCGTCAGATGTGTATAAGAGACAGACTGAGGCAGGAGGATCACT |
| ct10_T8_In_R | GTCTCGTGGGCTCGGAGATGTGTATAAGAGACAGGCCCATGACCTCCGTCTATG |
| ct10_T9_In_F | TCGTCGGCAGCGTCAGATGTGTATAAGAGACAGTCCCGTTGAGAACCACTTGG |
| ct10_T9_In_R | GTCTCGTGGGCTCGGAGATGTGTATAAGAGACAGCAGGTTCCAGTTCTGCCTGT |
| **20-target mgRNA** | |
| ct20_T1_In_F | TCGTCGGCAGCGTCAGATGTGTATAAGAGACAGCTACAGCTACTCGGGAGGCT |
| ct20_T1_In_F | GTCTCGTGGGCTCGGAGATGTGTATAAGAGACAGTACTCAGGAACAGGTGCGTG |
| ct20_T2_In_F | TCGTCGGCAGCGTCAGATGTGTATAAGAGACAGTAGCTGGTGTGGTGACATGC |
| ct20_T2_In_R | GTCTCGTGGGCTCGGAGATGTGTATAAGAGACAGCTGTCCCCTGTTAACACCCC |
| ct20_T3_In_F | TCGTCGGCAGCGTCAGATGTGTATAAGAGACAGAAAAATGGGCTTGGCTGGAC |
| ct20_T3_In_R | GTCTCGTGGGCTCGGAGATGTGTATAAGAGACAGCTGTACACAGCTAGGCCTGT |
| ct20_T4_In_F | TCGTCGGCAGCGTCAGATGTGTATAAGAGACAGTCCCAGCTACTTGAGAGGCT |
| ct20_T4_In_R | GTCTCGTGGGCTCGGAGATGTGTATAAGAGACAGCTCAGTAAGGGTGAGTTCTGGT |
| ct20_T5_In_F | TCGTCGGCAGCGTCAGATGTGTATAAGAGACAGCGCCTGTAATCCCAGCTACT |
| ct20_T5_In_R | GTCTCGTGGGCTCGGAGATGTGTATAAGAGACAGTGTCATGCTCTGGGTTGAGG |
| ct20_T6_In_F | TCGTCGGCAGCGTCAGATGTGTATAAGAGACAGTGGTGGTGCATGCCTGTAAT |
| ct20_T6_In_R | GTCTCGTGGGCTCGGAGATGTGTATAAGAGACAGGTGTCAGGTTAGCTTTGCAGTG |
| ct20_T7_In_F | TCGTCGGCAGCGTCAGATGTGTATAAGAGACAGCAGTTACTCGGGAGGCTGAG |

| | |
|---|---|
| ct20_T7_In_R | GTCTCGTGGGCTCGGAGATGTGTATAAGAGACAGACCATCCTGCTTCTGCTGTC |
| ct20_T8_In_F | TCGTCGGCAGCGTCAGATGTGTATAAGAGACAGAAGCACCTGCAATCCCATCT |
| ct20_T8_In_R | GTCTCGTGGGCTCGGAGATGTGTATAAGAGACAGTCTTGGGATCAGTCTCTGGGT |
| ct20_T9_In_F | TCGTCGGCAGCGTCAGATGTGTATAAGAGACAGGGCGCCTGTAATTCCAGCTA |
| ct20_T9_In_R | GTCTCGTGGGCTCGGAGATGTGTATAAGAGACAGTTGACTCAGACACACAGGGC |
| ct20_T10_In_F | TCGTCGGCAGCGTCAGATGTGTATAAGAGACAGCACCTGTAGTCCCAGCTGAG |
| ct20_T10_In_R | GTCTCGTGGGCTCGGAGATGTGTATAAGAGACAGTGCTTTCGTTACCGGTGTCA |
| ct20_T11_In_F | TCGTCGGCAGCGTCAGATGTGTATAAGAGACAGTGCACCTGTAGTCCCAGCTA |
| ct20_T11_In_R | GTCTCGTGGGCTCGGAGATGTGTATAAGAGACAGTTACAGGTGTGAGCCACTGC |
| ct20_T12_In_F | TCGTCGGCAGCGTCAGATGTGTATAAGAGACAGGGTCCTTGCTACTTGGGAGG |
| ct20_T12_In_R | GTCTCGTGGGCTCGGAGATGTGTATAAGAGACAGACCTTCTCTGCCTGTCCCTA |

**Supplementary Table 8: Oligos for sequencing mgRNA targets. PCR-3**

| **Name** | **Sequence (5′ to 3′) Red marks the i7/i5 8 bp index sequence** |
|---|---|
| NGS_Index_F1 | AATGATACGGCGACCACCGAGATCTACACCTCTCTATTCGTCGGCAGCGTC |
| NGS_Index_F2 | AATGATACGGCGACCACCGAGATCTACACTATCCTCTTCGTCGGCAGCGTC |
| NGS_Index_F3 | AATGATACGGCGACCACCGAGATCTACACGTAAGGAGTCGTCGGCAGCGTC |
| NGS_Index_F4 | AATGATACGGCGACCACCGAGATCTACACACTGCATATCGTCGGCAGCGTC |
| NGS_Index_F5 | AATGATACGGCGACCACCGAGATCTACACAAGGAGTATCGTCGGCAGCGTC |
| NGS_Index_F6 | AATGATACGGCGACCACCGAGATCTACACCTAAGCCTTCGTCGGCAGCGTC |
| NGS_Index_F7 | AATGATACGGCGACCACCGAGATCTACACCGTCTAATTCGTCGGCAGCGTC |
| NGS_Index_F8 | AATGATACGGCGACCACCGAGATCTACACTCTCTCCGTCGTCGGCAGCGTC |
| NGS_Index_F9 | AATGATACGGCGACCACCGAGATCTACACTCGACTAGTCGTCGGCAGCGTC |
| NGS_Index_F10 | AATGATACGGCGACCACCGAGATCTACACTTCTAGCTTCGTCGGCAGCGTC |
| NGS_Index_F11 | AATGATACGGCGACCACCGAGATCTACACCCTAGAGTTCGTCGGCAGCGTC |
| NGS_Index_F12 | AATGATACGGCGACCACCGAGATCTACACGCGTAAGATCGTCGGCAGCGTC |
| NGS_Index_R1 | CAAGCAGAAGACGGCATACGAGATTCGCCTTAGTCTCGTGGGCTCGG |
| NGS_Index_R2 | CAAGCAGAAGACGGCATACGAGATCTAGTACGGTCTCGTGGGCTCGG |
| NGS_Index_R3 | CAAGCAGAAGACGGCATACGAGATTTCTGCCTGTCTCGTGGGCTCGG |
| NGS_Index_R4 | CAAGCAGAAGACGGCATACGAGATGCTCAGGAGTCTCGTGGGCTCGG |
| NGS_Index_R5 | CAAGCAGAAGACGGCATACGAGATAGGAGTCCGTCTCGTGGGCTCGG |
| NGS_Index_R6 | CAAGCAGAAGACGGCATACGAGATCATGCCTAGTCTCGTGGGCTCGG |
| NGS_Index_R7 | CAAGCAGAAGACGGCATACGAGATGTAGAGAGGTCTCGTGGGCTCGG |
| NGS_Index_R8 | CAAGCAGAAGACGGCATACGAGATCAGCCTCGGTCTCGTGGGCTCGG |
